# Large-scale Nanopore sequencing resolves species diversity and distribution of European Ulva

**DOI:** 10.64898/2026.11.17.767378

**Authors:** Luna M. van der Loos, Sofie D’hondt, Emma Coulembier Vandelannoote, Quinten Bafort, Griet Casteleyn, Pınar Akdoğan Şirin, Stefano Accoroni, Umair Ahsan, Gerli Albert, Francisco Arenas, Simona Armeli Minicante, Edlira Aruci, Georgios Bakratsas, Karina Balina, Francisco R. Barboza, Ieva Barda, Angela G. Bartolo, Inka Bartsch, Jakob F. M. Becker, Zahira Belattmania, Giulia Bellanti, Ricardo Bermejo, Fabrizio Bernardi Aubry, Anaëlle Bizien, Aurelie Blanfuné, Jonas Blomme, Michael Bo Rasmussen, Teis Boderskov, Kruno Bonačić, Nathalie Bourgougnon, Paul Brazier, Juliet Brodie, Annette Bruhn, Francis Bunker, Thomas Burel, Julie Camus, João Canning-Clode, Alexander Chemodanov, Gercende Courtois de Viçose, Javier Cremades, Ajit Sharadrao Dake, Thibaut de Bettignies, Bruno de Reviers, Nathalie De Somer, Gonzalo Del Olmo, Julian Di Giandomenico, Pilar Díaz-Tapia, Aires Miguel Duarte, Luisa Pia Düsedau, Hannah Scarlett Earp, Marco Faasse, Annalisa Falace, Stefania Finotto, Daniela Freitas, Daniela Gabriel, Régis Gallon, Sara García-Poza, Verónica García-Redondo, Sindri Gíslason, Franz Goecke, Alexander Goldberg, Juan Luis Gomez Pinchetti, Lior Guttman, Jack R. Hall, Peter Hantz, Svenja Heesch, Mathieu Helias, Laurie C. Hofmann, Lasse Hornbek Nielsen, Vivian Husa, Ljiljana Iveša, Charlotte L Jennings, Sergi Joher, Veronika Valt, Mart Karremans, Petros Katapodis, Manfred Kaufmann, Ioannis Kefalas, Abigail Kennedy, Maximilian Knips, Jessica Knoop, Maria Kosmidou, Jonne Kotta, Frithjof C. Küpper, Kristoffer Larsen-Ledet, Gwenaelle Le Blay, Line Le Gall, Vincent Le Garrec, Delphie Le Thoër, Kitte Linding Gerlich, Kate Lock, Ana Lokovšek, Félix L. Figueroa, Petra Lučić, Christine Maggs, Erik-jan Malta, Francesco Paolo Mancuso, Antonio Manghisi, Luisa Mangialajo, Anna Maria Mannino, Oana Alina Marin, Joana Micael, David Milla-Figueras, Davide Moccia, Cátia Monteiro, Pippa J Moore, Marina Morabito, Ignacio Moreu, Liam Morrison, Robert J. Mrowicki, Konstantinia Nakou, Yvonne Nemcova, Francesca Neri, África Núñez García De la Morena, Wiebe Nijland, Anne-Marin Nisumaa-Comeau, Louka Noyen, Kristiina Nurkse, Sotiris Orfanidis, Martina Orlando-Bonaca, Apostolos Papadimitriou, Evangelos Papadimitriou, Viviana Pasquini, Sauvann Paulino, Sofie Peeters, Viviana Peña, Carolina Pena-Martín, Rui Pereira, Frank Perk, Francesca Petrucci, Laura Piazzese, Addis Pierantonio, José Pintado, Emeline Poisson, Laura Prigge, Raquel Quintã, Céline Rebours, Maggie Reddy, Simon Redlich, Kelli Richardon, Esben Rimi Christiansen, Fabio Rindi, Pedro Rodrigues, Valerie J. Rodrigues, Conxi Rodríguez-Prieto, Florence Rousseau, Andrzej S. Rybak, Brahim Sabour, Marta Sansón, Susanne Schäfer, Muki Shpigel, Marco Sigovini, Clara Simon, Nathalie Simon, Kjersti Sjøtun, Stamatia Spyrou, Haralambos Stamatis, Willem Stock, Sylvia Strauss, Heather Sugden, Ergün Taşkın, Thierry Thibaut, Gwendal Toullec, Eva Troianou, Ana Tronholm, Soultana Tsioli, Inci Tuney, Ioannis Tzovenis, Marika Ubaldi, José Utge, Lauren Vapillon, Vasily Vishnyakov, Sofie Vranken, Andrzej M. Woyda-Ploszczyca, Ante Žuljević, Miek Zwamborn, Ronan Sulpice, Thomas Wichard, Aschwin Engelen, Sophie Steinhagen, Frederik Leliaert, Olivier De Clerck

**Affiliations:** Phycology Research Group, Department of Biology, Ghent University, Ghent, Belgium; Naturalis Biodiversity Center, Leiden, Netherlands; Department of Plant Biotechnology and Bioinformatics, Ghent University, (Technologiepark 71,) 9052 Ghent, Belgium; VIB Center for Plant Systems Biology, VIB, (Technologiepark 71,) 9052 Ghent, Belgium; Fatsa Faculty of Marine Science, Department of Fisheries Technology Engineering, Ordu University, 52400, Fatsa, Ordu, Türkiye; Department of Life and Environmental Sciences, Università Politecnica delle Marche, Italy; Department of Plant and Animal Production, Burdur Vocational School of Food, Agriculture and Livestock, Burdur Mehmet Akif Ersoy University, Türkiye; Estonian Marine Institute, University of Tartu, Mäealuse 14, 12618, Tallinn, Estonia; CIIMAR -Interdisciplinary Centre of Marine and Environmental Research, FCUP -Faculty of Science, University of Porto, Portugal; Consiglio Nazionale delle Ricerche, Istituto di Scienze Marine (CNR-ISMAR), Venice, Italy; Department of Biotechnologies, Chemistry and Pharmacy, University of Siena, Siena, Italy; Laboratory of Biotechnology, Department of Biological Applications and Technologies, University of Ioannina, Ioannina, Greece; Institute of Microbiology and Biotechnology, University of Latvia, Jelgavas iela 1, Riga, Latvia; Latvian Institute of Aquatic Ecology, Agency of Daugavpils University, Riga, Latvia; Institute of Earth Systems, University of Malta, Malta; Alfred-Wegener Institute, Helmholtz Centre for Polar and Marine Research, Am Handelshafen 12, 27570 Bremerhaven, Germany; Phycology, Blue Biodiversity and Biotechnology RU, Laboratory of Plant Biotechnology, Ecology and Ecosystem Valorization – CNRST Labeled Research Unit N°10, Faculty of Sciences, University Chouaib Doukkali, El Jadida, Morocco; National Biodiversity Future Center, Palermo, 60131, Italy; Department of Ecology and Geology, University of Malaga, Bulevar Pasteur, 31, s/n, 29010 Málaga, Spain; Université Brest, CNRS, IRD, Ifremer, LEMAR, IUEM, F-29280 Plouzané, France; Aix Marseille University, University of Toulon, CNRS, IRD, MIO (Mediterranean Institute of Oceanography), Marseille, France; Department of Plant Biotechnology and Bioinformatics, Ghent University, Ghent, Belgium; VIB-UGent Center for Plant Systems Biology, Ghent, Belgium; Aarhus University, C.F. Møllers Allé 3, 8000 Aarhus, Denmark; Department of applied ecology, University of Dubrovnik, Croatia; Laboratoire de Biotechnologie et Chimie Marines, Université Bretagne Sud, EMR CNRS 6076, IUEM, 56000 Vannes, France; Natural Resources Wales, Maes Y Ffynnon, Ffordd Penrhos, Bangor, Gwynedd, LL57 2DW, United Kingdom; Natural History Museum, Research, Cromwell Road, London SW7 5BD, United Kingdom; MarineSeen, Estuary Cottage, Bentlass, Hundleton, Pembrokeshire SA71 5RN, United Kingdom; Univ Brest, CNRS, IRD, Ifremer, LEMAR, F-29280 Plouzané, France; Laboratoire des Sciences Appliquées de Cherbourg, UNICAEN, Normandie Université, France; Conservatoire National des Arts et Métiers, INTECHMER, France; MARE -Marine and Environmental Sciences Centre, ARNET – Aquatic Research Network; Agência Regional para o Desenvolvimento da Investigação Tecnologia e Inovação (ARDITI), Madeira, Portugal; Faculty of Life Sciences, University of Madeira, 9020-105 Funchal, Madeira, Portugal; Tel Aviv University, School of Mechanical Engineering, Tel Aviv, Israel; IU-ECOAQUA, GIA, Universidad de Las Palmas de Gran Canaria, Las Palmas, Spain; BioCost Research Group, Facultade de Ciencias and Centro Interdisciplinar de Química e Bioloxía (CICA), Universdade da Coruña, A Coruña, Spain; Marine Biology and Biotechnology Program, Department of Life Sciences, Ben-Gurion University of the Negev, Eilat Campus, Hatmarim Blvd. 162, Eilat/POB 653, Beer-Sheva, 8410501, Israel; PatriNat (OFB, MNHN), Muséum national d’Histoire naturelle, Paris, France; Centre de Recherche sur la Biodiversité et l’Environnement, Université de Toulouse, Toulouse INP, CNRS, IRD, Toulouse, France; Institut Systématique Evolution Biodiversité (ISYEB), Muséum national d’Histoire Naturelle, CNRS, Sorbonne Université, EPHE, Paris, France; ANEMOON Foundation, Netherlands; Institute of Marine Research (IIM-CSIC), Spain; CRETUS, Bioaplic Research Group, Universidade de Santiago de Compostela, Spain; The Dove Marine Laboratory, School of Natural and Environmental Sciences, Newcastle University, Newcastle upon Tyne, NE1 7RU, United Kingdom; Institute of Marine Research, His 4817, Norway; Eurofins AquaSense, Korringaweg 7, 4401NT, Yerseke, The Netherlands; Department of Life Sciences, University of Trieste, Italy; Greenaqua Lda. Rua Dr João Graça S/N. EN109 Km 66.2. 3840-403, Vagos, Portugal; CIBIO -Research centre in Biodiversity and genetic resources, Portugal; University of the Azores, Rua da Mãe de Deus, 9501-801 Ponta Delgada, Portugal; Departamento de Biotecnología-Instituto Tecnológico de Canarias, Spain; BioCost Research Group, Facultad de Ciencias and Centro Interdisciplinar de Química e Bioloxía (CICA), University of A Coruña, A Coruña, Spain; Southwest Iceland Nature Research Centre (SINRC), 245 Sudurnesjabær, Iceland; Norwegian University of Life Sciences (NMBU), Norway; Spanish Bank of Algae, Institute of Oceanography and Global Change, University of Las Palmas de Gran Canaria, Spain; The Department of Blue Biotechnologies and Sustainable Mariculture, Leon H. Charney School of Marine Science, University of Haifa, Haifa, Israel; Eötvös Loránd University, Dept. of Organic Chemistry, Budapest, Hungary; Fibervar Llc. Cluj/Kolozsvár, Romania; Applied Ecology & Phycology, Institute for Biological Sciences, University of Rostock, Albert-Einstein-Strasse 3, 18059 Rostock, Germany; Bremerhaven University of Applied Sciences, Germany; Pure Algae, Denmark; Institute of Marine Research, Norway; Ruđer Bošković Institute, Center for Marine Research, Rovinj, Croatia; Centre Oceanogràfic de les Balears, Instituto Español de Oceanografía (COB-IEO), CSIC. Parc Bit, Complex Balear de Recerca. Thomas Edison 5, planta baixa. 07121 Palma, Illes Balears, Spain; Department of Botany, Charles University, Prague, Czech Republic; Faculty of Life Sciences, University of Madeira, Portugal; MARE – Marine and Environmental Sciences Centre / ARNET -Aquatic Research Network, Madeira, Portugal; Fisheries Research Institute (ELGO-DIMITRA), Nea Peramos, Kavala, Greece; University of Aberdeen, Scotland, United Kingdom; Friedrich Schiller University Jena, Institute for Inorganic and Analytical Chemistry, Germany; Department of Estuarine and Delta Systems, Royal Netherlands Institute for Sea Research (NIOZ), Yerseke, Netherlands; Marine Biodiscovery Centre, Department of Chemistry, University of Aberdeen, Aberdeen AB24 3UE, Scotland, United Kingdom; Department of Biology, San Diego State University, California, CA 92182-1030, USA; Oceanography Center, University of Cyprus, 1 Panepistimiou Av., 2109 Aglandjia, Nicosia, Cyprus; Station Biologique de Roscoff, Sorbonne University and CNRS, Roscoff, France; Univ Brest, CNRS, IRD, Observatoire, IUEM, F-29280 Plouzané, France; Cellule de Suivi du Littoral Normand (CSLN), France; Natural Resources Wales, Skomer MCZ, Martins Haven, Pembrokeshire Wales SA623BJ, United Kingdom; Marine Biology Station Piran, National Institute of Biology, Slovenia; University of Malaga, Instituto Andaluz de Biotecnología y Desarrollo Azul (IBYDA), Loma de San Julián, 2, 29004 Malaga; Institute of Oceanography and Fisheries, Split, Croatia; School of Biological Sciences, Queen’s University Belfast, 19 Chlorine Gardens, Belfast BT9 5DL, Northern Ireland, United Kingdom; Aquaculture Technology Centre, CTAQUA, Muelle Comcercial S/N, 11500, El Puerto de Santa Maria, Spain; Department of Earth and Marine Sciences (DiSTeM), Palermo, Italy; University of Messina, Italy; Université Côte d’Azur, CNRS, ECOSEAS, France; Department of Biological, Chemical and Pharmaceutical Sciences and Technologies (STeBiCeF), Palermo, Italy; National Institute for Marine Research and Development “Grigore Antipa” (NIMRD), Constanta, Romania; University of the Azores, Institute of Marine Sciences -OKEANOS, Rua Professor Doutor Frederico Machado 4, 9900-140 Horta, Portugal; Department of Life and Environmental Sciences, University of Cagliari, Cagliari, Italy; BIOPOLIS Program in Genomics, Biodiversity and Land Planning; University of Malaga, Instituto Andaluz de Biotecnología y Desarrollo Azul (IBYDA), Bulevar Pasteur, 31, s/n, 29010 Málaga, Spain; School of Natural Sciences, Ryan Institute, University of Galway, Ireland; Department of Ecology and Geology, University of Malaga, Bulevar Pasteur, 31, s/n, 29071 Málaga, Spain; Department of Physical Geography, Utrecht University, Netherlands; Departemento Ciencies Ambientals i Recursos Naturals. Universitat d’Alacant, Spain; A4F, Algafuel. Estrada Paço do Lumiar, Campus do Lumiar Ed E R/C. 1649-038, Lisbon, Portugal; MARE -Marine and Environmental Sciences Centre, ARNET – Aquatic Research Network, Agência Regional para o Desenvolvimento da Investigação, Tecnologia e Inovação (ARDITI), 9020-105 Funchal, Madeira, Portugal; Institute of Marine Research (IIM-CSIC), Vigo, Galicia, Spain; University of Porto, Faculty of Sciences, Department of Biology, Portugal; S2AQUA -Collaborative Laboratory, Association for a Sustainable and Smart Aquaculture, Portugal; Møreforsking AS, Borgundveien 340, 6009 Ålesund, Norway; School of Chemical and Biological Sciences, Ryan Institute, University of Galway, Ireland; Dipartimento di Scienze della Vita e dell’Ambiente, Università Politecnica delle Marche, Ancona, Italy; Rif Field Station, 675 Raufarhöfn, Iceland; Natural Science Institute of Iceland, 300 Akranes, Iceland; Institute of Biological, Environmental and Rural Sciences, Aberystwyth University, Gogerddan, Aberystwyth, SY23 3EE, United Kingdom; Universitat de Girona. Facultat de Ciències. Ma Aurèlia Capmany 69. 17003, Girona, Spain; Department of Hydrobiology, Institute of Environmental Biology, Faculty of Biology, Adam Mickiewicz University, Poznan, Uniwersytetu Poznanskiego 6, PL 61-614 Poznan, Poland; Departamento Botánica, Ecología y Fisiología Vegetal, Universidad de La Laguna, Canary Islands, Spain; The Leon H. Carney School of Marine Sciences, University of Haifa, Israel; Department of Biological Sciences, University of Bergen, Bergen, Norway; The Seaweed Company, Netherlands; Department of Biology, Faculty of Engineering and Natural Sciences, Manisa Celal Bayar University, Manisa, Türkiye; Technologique et Scientifique de Luminy, 163 avenue de Luminy – Bâtiment Méditerranée, 13288 Marseille cedex 09, France; Kefalonia Fisheries SA, Greece; Department of Biological and Environmental Sciences, University of Gothenburg, Box 463, 405 30 Göteborg, Sweden; Gothenburg Global Biodiversity Centre, Box 463, 413 90 Göteborg, Sweden; Ege University, Faculty of Science, Department of Biology, Bornova-Izmir, Türkiye; Papanin Institute for Biology of Inland Waters, Russian Academy of Sciences, Borok, Yaroslavl, Russia; Department of Bioenergetics, Institute of Molecular Biology and Biotechnology, Faculty of Biology, Adam Mickiewicz University, Poznan, Uniwersytetu Poznanskiego 6, PL 61-614 Poznan, Poland; Knockvologan Studio, Isle of Mull, United Kingdom; Centro de Ciências do Mar do Algarve (CCMAR/CIMAR LA), Campus de Gambelas, Universidade do Algarve, 8005-139 Faro, Portugal; University of Algarve, Portugal; Department of Marine Sciences-Tjärnö, University of Gothenburg, SE-452 96 Strömstad, Sweden; Department of Natural History, University Museum, University of Bergen, P.O. Box 7800, 5020 Bergen, Norway; Meise Botanic Garden, Nieuwelaan 38, B-1860 Meise, Belgium

**Keywords:** biodiversity, biogeography, Ulva, high-throughput barcoding

## Abstract

The green algal genus Ulva is widespread and ecologically important in European coastal ecosystems. However, species diversity and distributions remain poorly resolved due to phenotypic plasticity and historical taxonomic uncertainty. By adopting a large-scale sampling and barcoding effort, we provide a unique continent-scale assessment of Ulva species diversity and distribution. We developed a novel high-throughput Oxford Nanopore Technologies sequencing strategy, pooling 768 samples (eight 96-well plates) per run and sequencing tufA and ITS simultaneously. Nanopore barcoding was accurate and efficient, with 96.3% of samples yielding at least one informative marker. Consensus sequences showed 99.4–100% similarity to Sanger sequences. By analysing 3,657 samples from 30 countries, we identified 46 species with contrasting distribution patterns and substantially expanded their known geographic ranges. This includes numerous country-level first records and the first records of Ulva howensis outside Australia and U. meridionalis within the study area. Five non-indigenous Ulva species were also detected, including widespread occurrences of U. australis. However, 10 species could not be assigned to named taxa because suitable reference sequences were unavailable or unreliable, highlighting persistent taxonomic gaps in public databases. Our findings demonstrate the power of coordinated, high-throughput molecular surveys to resolve cryptic marine biogeography at continent scales. The resulting high-resolution spatial baseline facilitates the early detection of non-indigenous species and range expansions. It also enables coastal managers to monitor invasion pathways, evaluate ecosystem risks associated with opportunistic bloom-forming taxa, and detect climate-driven distribution shifts across European waters.

## 1. Introduction

The green algal genus Ulva (Ulvophyceae, Chlorophyta) is among the most widespread and ecologically important macroalgae. It occurs abundantly across a broad range of aquatic environments, including marine, brackish, and freshwater ecosystems from Arctic to tropical regions. Species of Ulva contribute substantially to primary production and nutrient cycling in the ecosystems where they occur. Their ability to thrive under variable salinity, temperature, and light conditions contributes to their success across diverse habitats (Messyasz & Rybak, 2011; Poole & Raven, 1997). Under nutrient-rich conditions, this ecological flexibility can result in prolific growth and the development of extensive green tides (Gao et al., 2010; Wan et al., 2017), with significant ecological and socioeconomic consequences, including alterations of ecosystem functioning, reduced biodiversity, deterioration of water quality, and negative impacts on fisheries, aquaculture, and coastal tourism (Smetacek & Zingone, 2013; Ye et al., 2011). In addition to its ecological importance, the high-growth capacity of Ulva has attracted considerable economic interest for applications in aquaculture, animal feed, human nutrition, wastewater remediation, and the extraction of bioactive compounds for the nutraceutical and cosmetic industries (Hofmann et al., 2025; Wichard et al., 2026).

Despite its widespread occurrence in marine ecosystems and growing economic importance, resolving species diversity within Ulva remains challenging due to its simple morphology, extensive morphological plasticity, and widespread phenotypic convergence (Lamb et al., 2025). Environmental factors such as light availability, nutrient concentrations, salinity, hydrodynamic conditions, and life-history stage can strongly influence thallus morphology and cytological characteristics (e.g., plastid shape and position, number of pyrenoids), causing individuals of the same species to exhibit markedly different forms (Blomster et al., 2002; Steinhagen et al., 2019). Conversely, distinct species may display virtually identical morphologies. As a result, traditional taxonomic approaches based on morphological characteristics have frequently led to misidentifications, incorrect assessments of regional biodiversity, and obscured biogeographic and diversity patterns. This issue is further compounded by the presence of phylogenetically distant green algal “look-alikes” (e.g., members of the Kornmanniaceae and Ulotrichales), which are regularly misidentified as Ulva in the field. Although 101 species of Ulva are currently formally described (Guiry & Guiry, 2026), morphological characters alone are often insufficient for reliable species identification, highlighting the need for molecular analyses to resolve species boundaries and distinguish closely related taxa.

While several molecular surveys of Ulva have been conducted (Supplementary Table S1), these studies remain geographically fragmented, typically focusing on individual countries, regions, or habitat types. In addition, different DNA barcode markers have been used across studies, hampering direct comparisons of species diversity, abundance, distributions, and ecological preference at the European scale. In order to overcome these shortcomings, we conducted the first continent-wide molecular assessment of Ulva diversity in Europe, using a standardised DNA barcoding panel consisting of the internal transcribed spacer of the ribosomal cistron (ITS) and elongation factor Tu (tufA). Through a coordinated consortium effort, we sampled Ulva specimens, along with morphologically similar bladed and tubular taxa from Ulvaceae, Kornmanniaceae, and Ulotrichales, across Europe and adjacent regions, spanning from the Black Sea to Macaronesia along an east–west axis and from Svalbard to Morocco along a north–south axis.

Sanger sequencing of multiple markers across thousands of specimens is restricted by high operational costs and labour-intensive workflows. To address this bottleneck, we developed and validated a high-throughput DNA barcoding pipeline based on Oxford Nanopore Technologies (ONT) that enabled us to process a large number of samples in an efficient and standardised manner. We used this improved taxonomic framework to reassess the diversity and distribution of Ulva across Europe. Because many historical records were based solely on morphological identifications, their reliability is questionable in light of current taxonomic knowledge.

By combining newly collected specimens with existing molecular data from across Europe, we aimed to develop a comprehensive continental-scale assessment of Ulva species diversity, clarify distribution patterns, and improve our understanding of the genus’ biogeography throughout Europe.

## 2. Methods

### 2.1 Study area and sample collection

Emerging within the context of the SeaWheat COST Action (Buck & Shpigel, 2023), the European Ulva Taxonomy Initiative (EULVA) consortium was established by contacting a broad network of phycologists across Europe, inviting them to collect Ulva specimens along their respective coastlines (https://sites.google.com/view/eulva) (Fig. 1). To ensure maximum consistency, all participating partners adhered to a standardized protocol for specimen collection, ecological data recording, and vouchering, including silica-gel tissue preservation for molecular analyses, field photography, and the preparation of herbarium vouchers where possible. Through this coordinated, collaborative framework, a total of 3,657 specimens were collected and sequenced (Supplementary Table S2). Target species included foliose and tubular members of the Ulvaceae (Percursaria, Ulva, Ulvaria, Umbraulva), but also species closely resembling Ulva in morphology, including members of the Kornmanniaceae (Blidingia, Kornmannia) and Ulotrichales (Gayralia, Monostroma, Protomonostroma, Pseudothrix). The study area comprised the Arctic Ocean, the Northeast Atlantic Ocean (including the North Sea, Baltic Sea, and Macaronesia), the Mediterranean Sea, and the Black Sea. Samples were collected in 30 countries, spanning from Svalbard in the north to Morocco in the south, and from the Azores in the west to Azerbaijan in the east (Fig. 2). To maximize species discovery, a wide range of habitats were investigated, including marine rocky reefs, maerl beds, intertidal mudflats, harbours, artificial breakwaters, coastal lagoons, salt marshes, and freshwater and brackish environments (Fig. 3). From each specimen, a subsample of ∼1cm^2^ tissue was cleaned gently with a paper towel, blotted dry and conserved in silica for subsequent molecular analyses. Where possible, specimens were also photographed, and the remaining thalli were preserved as herbarium vouchers and deposited in herbarium collections in their country of origin (Supplementary Table S2).

**Figure 1.**
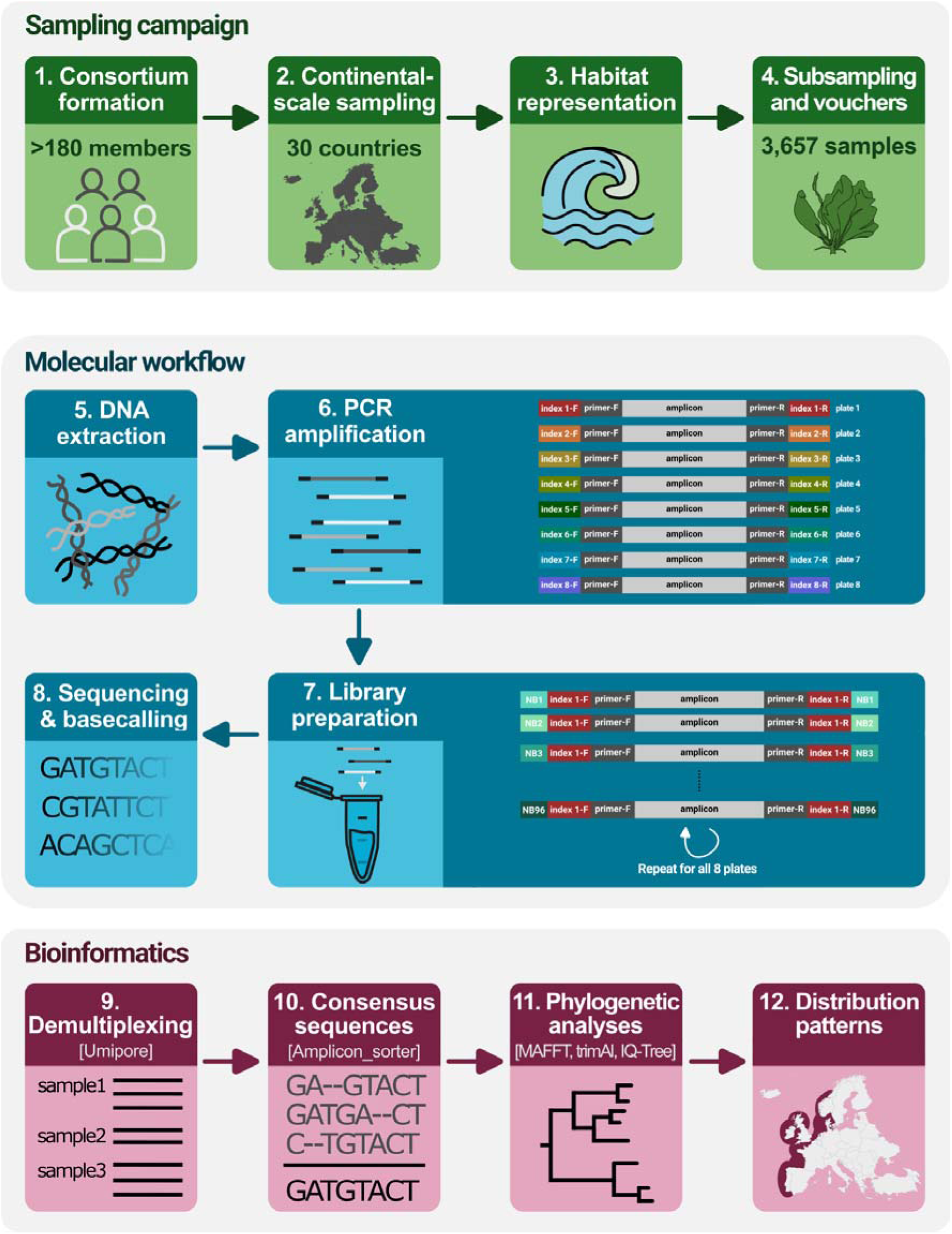
Overview of the EULVA workflow, from continental-scale sampling through DNA sequencing and bioinformatic processing of 768 samples (eight 96-well plates) using Oxford Nanopore Technologies.

**Figure 2.**
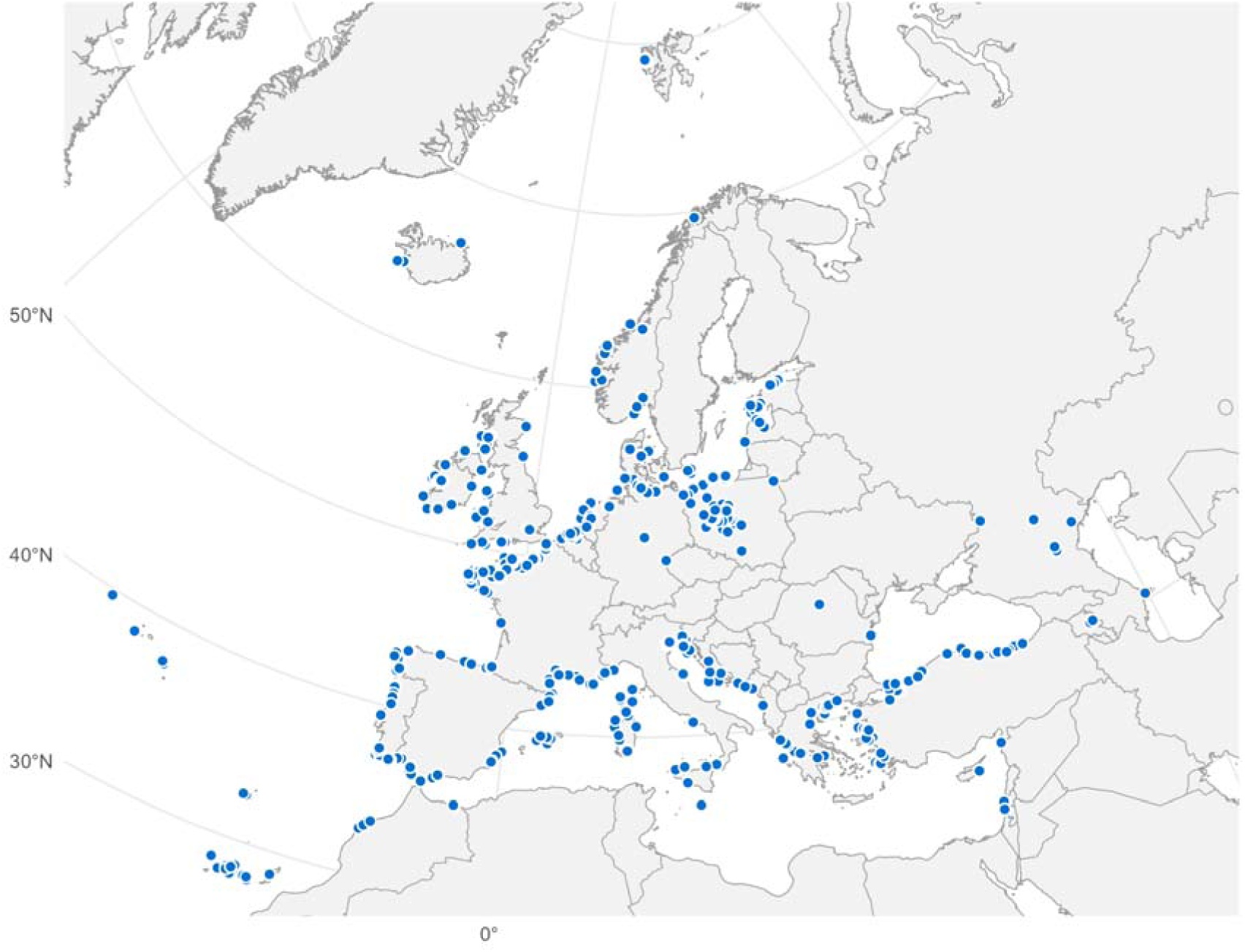
Map of the study area showing the collection localities of the 3,657 samples analysed by the EULVA consortium.

**Figure 3.**
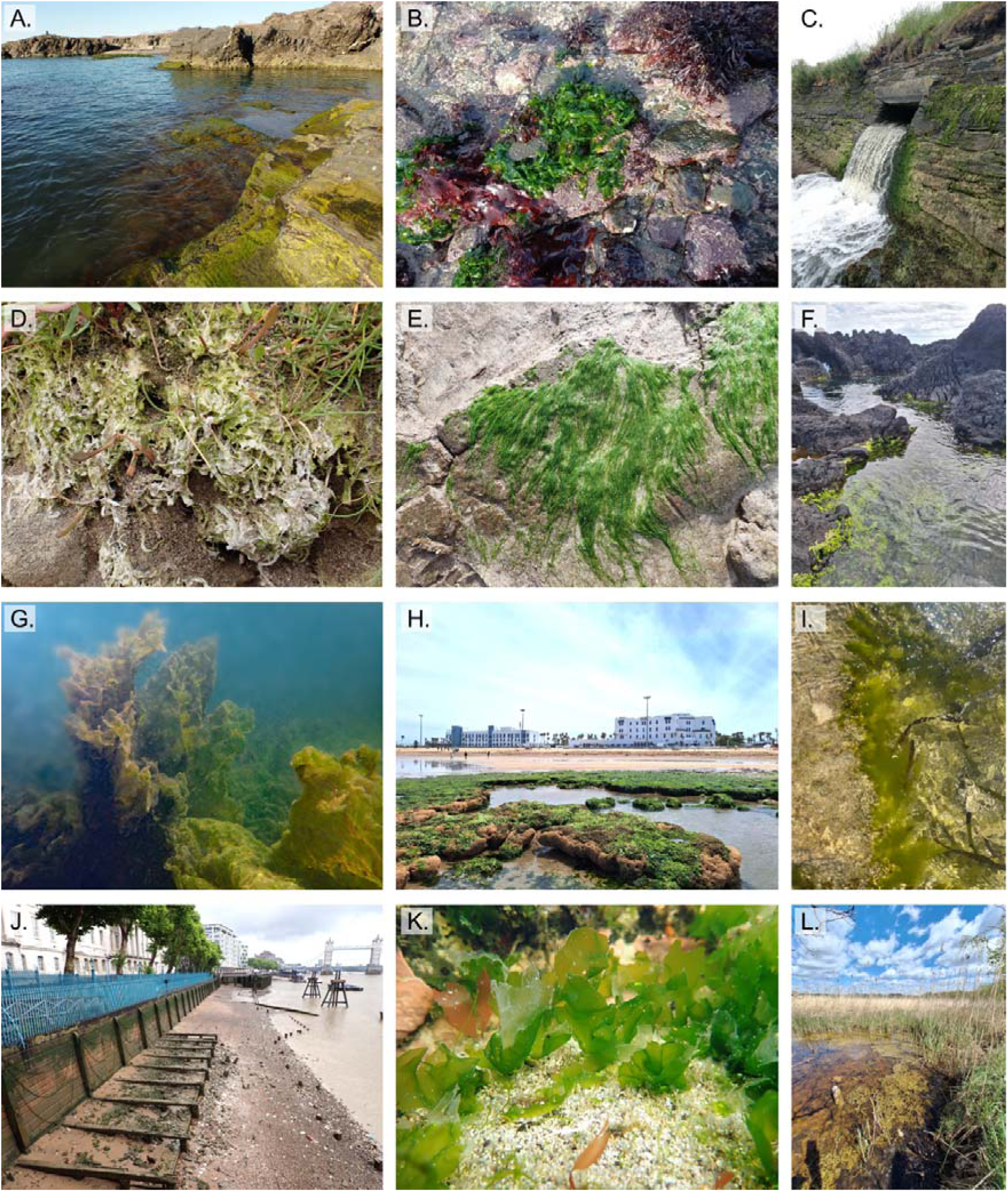
Examples of the various habitats sampled. (A) Pseudothrix borealis and Kornmannia species in the Arctic Ocean. Kapp Guissez, Svalbard (Photo by Luisa Düsedau) (B) Ulva lacinulata on the intertidal of the rocky Atlantic coast. Brittany, France (Photo by Anaëlle Bizien) (C) Blidingia marginata near a water outlet. Brittany, France (Photo by Thomas Burel) (D) Ulva intestinalis in a coastal saltmarsh. Wales, UK (Photo by Paul Brazier) (E) Ulva prolifera on a clay bank in a tidal mudflat. Wadden Sea, Netherlands (Photo by Luna van der Loos) (F) Ulva compressa on volcanic rock. Azores, Macaronesia (Photo by Yvonne Nemcova) (G) Ulva australis in the subtidal. Marseille-Rhône canal, France (Photo by Marco Faasse) (H) Ulva lactuca on a rocky plateau. El Jadida, Morocco (Photo by Zahira Belattmania) (I) Ulva tepida on the Mediterranean coast. St. Paul’s Bay, Malta (Photo by Angela G. Bartolo) (J) Ulva intestinalis and Blidingia cornuta along the Thames river. London, UK (Photo by Luna van der Loos) (K) Ulvaria splendens in a tidal pool. Melrakkaslétta, Iceland (Photo by Joana Micael). (L) Ulva sp. 5 sensu Steinhagen in a landlocked lake/saltmarsh. Soos, Czech Republic (Photo by Yvonne Nemcova).

### 2.2 Molecular analyses

To process the large number of samples, we developed a high-throughput sequencing strategy using Oxford Nanopore Technologies, in which 768 samples (eight 96-well plates) were pooled and sequenced using dual markers (tufA and ITS) in a single sequencing run (Fig. 1). These markers have been shown to capture Ulva diversity and are the most commonly used molecular markers for the genus (Tran et al., 2022). For each marker, our approach combined eight indexed primer pairs with 96 unique Nanopore native barcodes ligated during the library preparation. All 96 samples within a given well plate were amplified using the same indexed primer pair and subsequently received unique Native Barcodes. In total, eight sequencing runs were performed to process all samples, using two runs per single flow cell.

Genomic DNA was extracted from silica-preserved samples in the molecular lab at Ghent University using a CTAB protocol (Doyle & Doyle, 1987; van der Loos et al., 2025). The chloroplast-encoded tufA gene and the internal transcribed spacer (ITS) were amplified with indexed primer pairs (Supplementary Table S3), based on primers tufAF/tufAR (Verbruggen et al., 2009) and ITS1/ITS4 (Mareš et al., 2011), respectively, each extended with one of the eight indexes. PCR mixtures consisted of 7.5 μl Phire Tissue Direct PCR Master Mix, 0.3 μl 50 x primer mix, 6.2 μl water, and 1.0 μl template DNA. TufA PCRs were run with a thermal profile of 3 min initial denaturation at 98°C, 40 cycles (98°C 10 s, 45°C 30 s, 72°C 30 s), and final extension of 7 min at 72°C. ITS PCRs were run with a thermal profile of 3 min initial denaturation at 98°C, 35 cycles (98°C 8s, 52°C 8s, 72°C 30 s), and final extension of 3 min at 72°C.

The sequencing libraries were prepared with the SQK-NBD114.96 kit according to the manufacturer’s protocol (Oxford Nanopore Technologies). After the native barcode ligation step was completed, the eight plates of samples were pooled and sequenced on a MinION with an R10.4 flow cell for 14-24 h (until 16,000 reads per barcode were generated).

For samples that could not be successfully sequenced using the high-throughput Nanopore approach described above, or that were received after completion of the Nanopore sequencing runs, tufA sequences were obtained using Sanger sequencing (n = 156). Methods followed van der Loos et al. (2025) and sequencing was performed by Macrogen (Amsterdam, the Netherlands). Samples identified as Kornmannia spp. did not amplify using the standard tufA primers and the tufA fragment was instead amplified with newly developed primers Kornmannia-F (5’-AACGACAGCGCGCCATTACG-3’) and Kornmannia-R (5’-CTGTGCTTCAAATCGGTTATGAGG-3’).

### 2.3 Bioinformatics

Raw reads were basecalled with MinKNOW v24.11.8 (in super-accurate basecalling mode v4.3.0, 400 bps) and filtered on a minimum Q-score of 10 and a minimum length of 400 bp. Data were subsequently demultiplexed with Umipore (Vierstraete, 2026) using the one-sided barcode setting and the default error percentage (15%) (Fig. 1). Consensus sequences were generated with Amplicon_sorter (Vierstraete & Braeckman, 2022) based on a similarity percentage of 98%, using 2,000 reads randomly selected per sample. Consensus sequences based on at least 100 reads were retained, aligned with MAFFT v.7.490 (Katoh et al., 2002), and trimmed with trimAl v1.4.rev15 using gappyout settings (Capella-Gutiérrez et al., 2009). Maximum likelihood phylogenetic trees were generated with IQ-Tree v2.2.0 using an automated model finder with 1,000 ultrafast bootstrap iterations (Nguyen et al., 2015). Sequences were identified based on peer-reviewed reference sequences retrieved from NCBI GenBank, including all relevant species from the study area and currently available type sequences.

Data were processed and visualised in R (R Core Team, 2025) using the _DPLYR_ (Wickham et al., 2026), _GGPLOT_2 (Wickham, 2016), _GRID_E_XTRA_ (Auguie, 2010), _PURRR_ (Wickham & Henry, 2015), _SCALES_ (Wickham et al., 2011), _SF_ (Pebesma, 2018; Pebesma & Bivand, 2023), and _STRINGR_ (Wickham, 2009) packages.

### 2.4 Benchmarking with Sanger-derived sequences

To benchmark the quality and accuracy of the Nanopore-based consensus sequences, we compared a subset with Sanger-based sequencing data derived from the same samples for both tufA and ITS regions. This set included 25 different species across 100 samples for tufA, and 13 different species across 39 samples for ITS. Percent identity matrices were generated with Clustal Omega (Sievers et al., 2011) using the --_PERCENT_-_ID_ option.

### 2.5 Voucher and data deposition

Physical voucher specimens prepared during this study were deposited in recognized herbaria within their respective countries of collection (Supplementary Table S2). For each specimen, collection data, images of fresh thalli or herbarium vouchers, and the DNA sequence data were archived in the Barcode of Life Data Systems (BOLD; https://www.boldsystems.org) under project EULVA with accession numbers EULVA001-26 to EULVA3679-26. In addition, sequences were deposited in GenBank (accession numbers XX-XX).

## 3. Results

### 3.1 Sequencing quality and success

To benchmark the quality of Nanopore-derived consensus sequences, we compared a subset of sequences generated from Nanopore data with Sanger-derived sequences obtained from the same samples. The ITS alignment included 13 species and was 573 bp in length. Pairwise genetic divergence across these species ranged from 64.0% to 98.4%. For all ITS samples (n = 39), Nanopore-based consensus sequences were identical or nearly identical (99.4 −100% similarity) to their Sanger-derived counterparts (Fig. 4). The 760 bp tufA alignment included 25 species, with pairwise genetic divergence ranging from 78.1% to 99.3%. For all tufA samples (n = 100), Nanopore-based consensus sequences were identical (100% similarity) to their Sanger-derived counterparts (Fig. S1).

**Figure 4.**
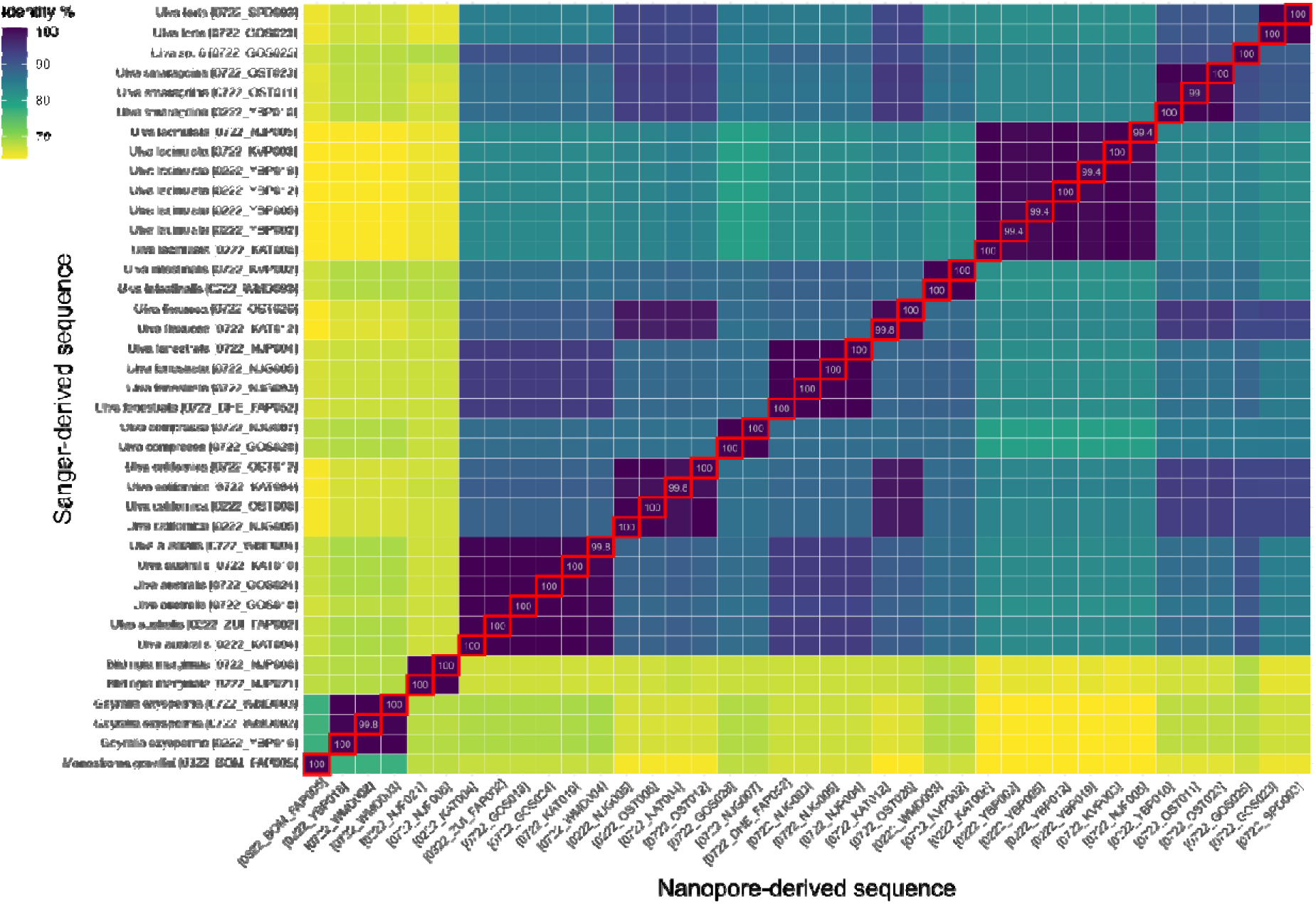
Pairwise percentage identity matrix of Sanger-and Nanopore-derived ITS sequences from 39 samples across 13 species. Colour represents percentage identity. Red-outlined tiles indicate comparisons between Sanger and Nanopore sequences originating from the same sample.

To further assess the effect of read depth on consensus accuracy, we compared consensus sequences generated from 100, 50, 25, and 10 randomly selected reads from the original FASTQ files. Across all tested samples (n = 50 for tufA and n = 39 for ITS), 25 reads were sufficient for Amplicon_sorter to generate consensus sequences that were 100% identical to those generated using higher read depths and to the corresponding Sanger sequences. Thus, reducing the read depth to 25 reads did not affect consensus accuracy in any of the tested samples. For the majority of samples, 10 reads were also sufficient to generate an accurate consensus sequence. However, no consensus sequence was generated from 10 reads in 11/39 ITS and 6/50 tufA samples. In a small number of additional cases, consensus sequences generated from 10 reads differed slightly from those generated at higher read depths, with 99.3–99.8% sequence identity.

Of the 3,636 samples originally submitted to the EULVA project for Nanopore sequencing, 3,501 (96.3%) yielded at least one informative sequence for either the tufA or ITS marker corresponding to a target species. For 30 samples, only non-target sequences were recovered (belonging, for example, to bacteria, fungi, or microalgae), while 105 samples produced an insufficient number of reads for reliable post-sequencing processing. Sequences for both tufA and ITS were successfully obtained from 2,875 samples (82.1%). An additional 457 samples (13.1%) yielded only an ITS sequence, whereas 169 samples (4.8%) yielded only a tufA sequence. This resulted in a total of 3,332 Nanopore-derived ITS sequences and 3,044 Nanopore-derived tufA sequences (3,200 tufA sequences when including Sanger-generated sequences).

### 3.2 Molecular species identification

A total of 46 species were identified across 3,657 samples (Fig. 5, Fig. 6). Among these, 17 species were identified using reference sequences derived from type specimens: Blidingia cornuta Steinhagen & Weinberger [holotype], B. minima (Nägeli ex Kützing) Kylin [epitype], Pseudothrix borealis Hanic & S.C.Lindstrom [holotype], Ulva australis Areschoug [holotype], U. californica Wille [lectotype], U. capillata Steinhagen [holotype], U. compressa Linnaeus [epitype], U. fenestrata Postels & Ruprecht [holotype], U. intestinalis Linnaeus [epitype], U. lacinulata Lightfoot [lectotype], U. lactuca Linnaeus [cross-referenced with rbcL sequence of the holotype], U. linza Linnaeus [epitype], U. meridionalis Horimoto & S.Shimada [holotype], U. ohnoi Hiraoka & S.Shimada [holotype], U. rigida C.Agardh [lectotype], U. smaragdina (Kützing) Hughey, Maggs, L.M.Loos, S.A.Harris & P.W.Gabrielson [lectotype], and U. tepida Masakiyo & S.Shimada [holotype].

**Figure 5.**
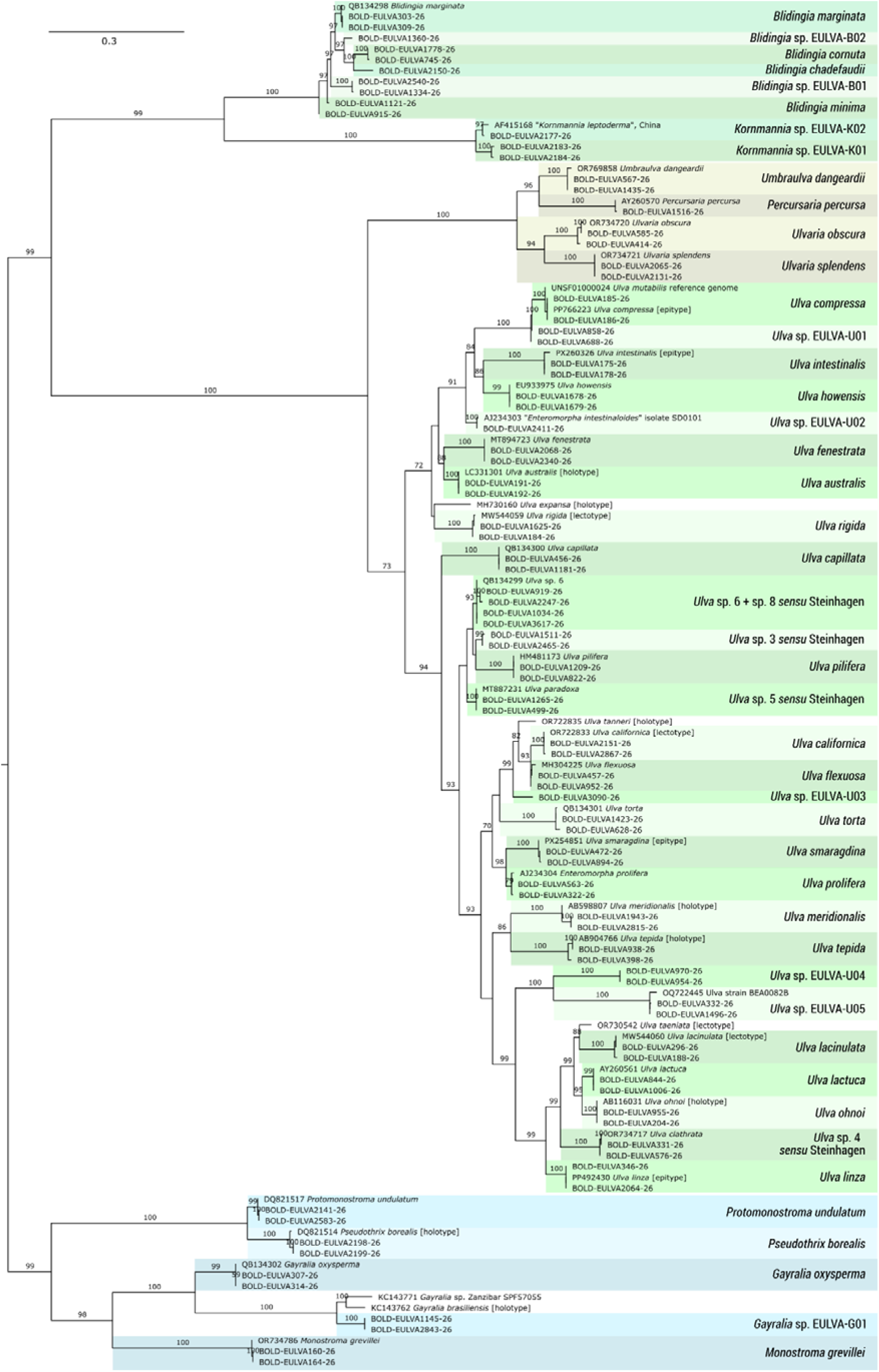
Maximum likelihood (IQ-TREE) phylogenetic tree of the 46 species identified within the EULVA dataset based on ITS sequences. Species found in this study are highlighted in colour. When available, two sequences per species from this study were included, with sequence accession numbers corresponding to the BOLD identifier. Representative sequences from GenBank for species relevant to the study area were included, as well as available Ulva type material for broader context. Branch support values represent ultrafast bootstrap support (UFBoot2 with 1,000 replicates). Branch lengths are scaled according to the number of nucleotide substitutions per site. See Suppl. Fig. S2 for a phylogenetic tree containing all specimens from this study.

**Figure 6.**
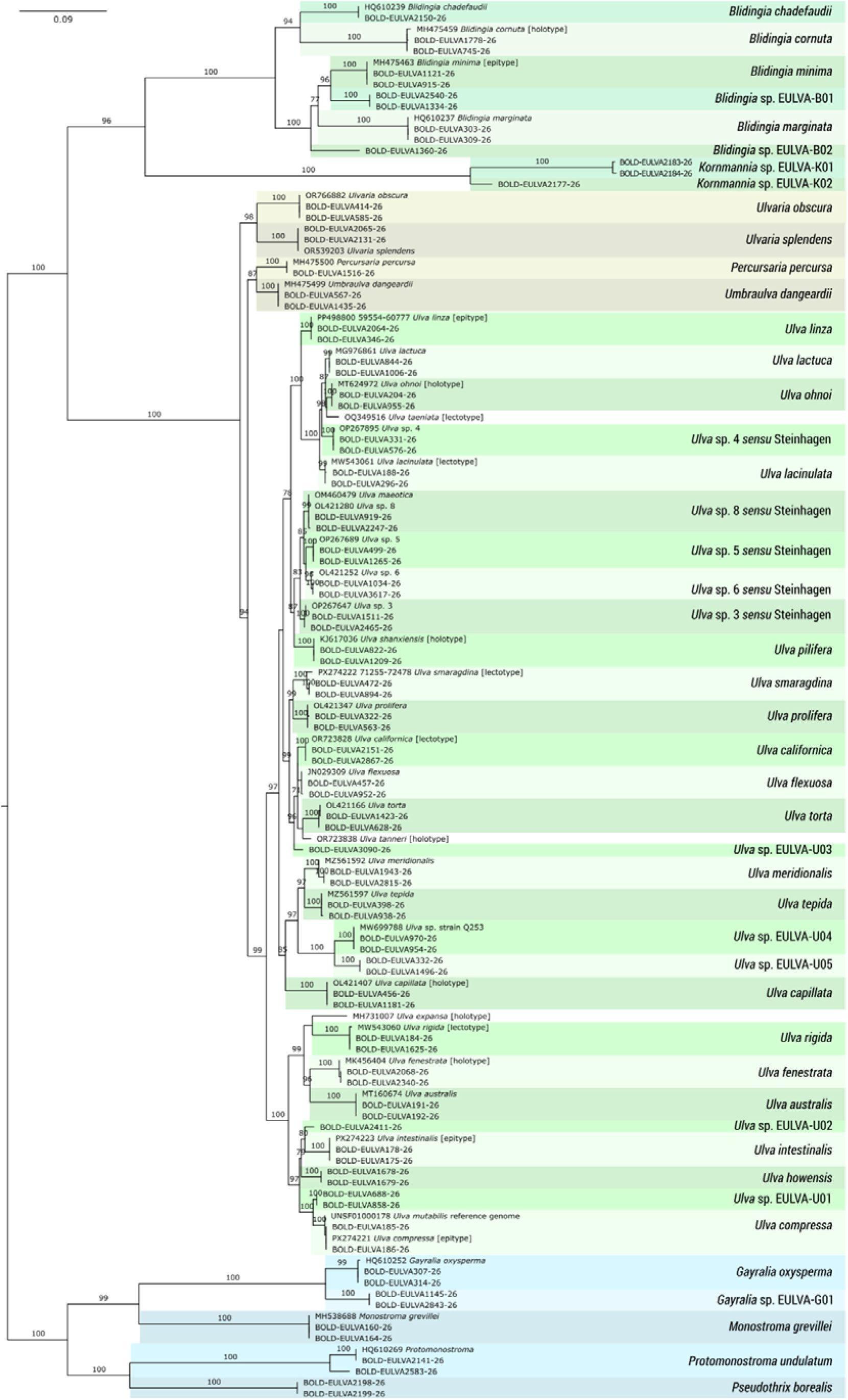
Maximum likelihood (IQ-TREE) phylogenetic tree of the 46 species identified within the EULVA dataset based on tufA gene sequences. Species found in this study are highlighted in colour. When available, two sequences per species from this study were included, with sequence accession numbers corresponding to the BOLD identifier and sample number in brackets. Representative sequences from GenBank for species relevant to the study area were included, as well as available Ulva type material for broader context. Branch support values represent ultrafast bootstrap support (UFBoot2 with 1,000 replicates). Branch lengths are scaled according to the number of nucleotide substitutions per site. See Suppl. Fig. S3 for a phylogenetic tree containing all specimens from this study.

In addition, 100 specimens – exclusively collected from freshwater habitats – matched the holotype of Ulva shanxiensis Le Chen, J.Feng & S.L.Xie based on the tufA marker, while their ITS sequences matched a reference sequence originally identified as Ulva flexuosa subsp. pilifera, now accepted as Ulva pilifera (Kützing) Škaloud & Leliaert.

For an additional 18 taxa, only non-type sequences were available in GenBank: Blidingia chadefaudii (Feldmann) Bliding, B. marginata (J.Agardh) P.J.L.Dangeard ex Bliding, Gayralia oxysperma (Kützing) K.L.Vinogradova ex Scagel & al., Monostroma grevillei (Thuret) Wittrock, Percursaria percursa (C.Agardh) Rosenvinge, Protomonostroma undulatum (Wittrock) K.L.Vinogradova, Ulvaria obscura (Kützing) Gayral ex Bliding, Ulvaria splendens (Ruprecht) K.L.Vinogradova, Umbraulva dangeardii M.J.Wynne & G.Furnari, Ulva flexuosa Wulfen, U. howensis (A.H.S.Lucas) Kraft, U. prolifera O.F.Müller, U. torta (Mertens) Trevisan, and Ulva sp. 3, sp. 4, sp. 5, sp. 6, and sp. 8 sensu Steinhagen et al. (2023). Among these, ITS sequences of Ulva sp. 5 (assigned based on tufA) matched GenBank accessions listed as Ulva paradoxa, and ITS sequences of Ulva sp. 4 (also assigned based on tufA) matched GenBank accessions identified as Ulva clathrata. Based on ITS only, Ulva sp. 6 and sp. 8 were difficult to differentiate.

Finally, 10 species were identified for which no matching sequences were available in public databases, or for which available matches were known to be incorrect. These unnamed species included two Blidingia species (here listed as Blidingia sp. EULVA-B01 and B02), a single Gayralia species (Gayralia sp. EULVA-G01), five Ulva species (Ulva sp. EULVA-U01 to U05), and two species tentatively assigned to the genus Kornmannia (Kornmannia sp. EULVA-K01 and K02). The Kornmannia clade, comprising 15 specimens collected in Iceland and Svalbard (Arctic), formed a sister clade to Blidingia.

Phylogenetic trees are shown in Figures 5 and 6, with each taxon represented by a reference sequence and two sequences from the EULVA dataset. The complete phylogenetic trees are provided in the Supplementary Materials.

### 3.3 Geographic distribution patterns and new records

The EULVA dataset revealed marked differences in the abundance, distribution, and habitat preferences of Ulvales and Ulotrichales species across the study area. While several species were widespread throughout the study area, others exhibited distinct biogeographic patterns or habitat preferences, with distributions predominantly restricted to Arctic, Atlantic, Mediterranean, or freshwater environments (Fig. 7). The dataset also substantially expands the known geographic ranges of several taxa, including numerous first records for individual countries and, in some cases, the first documented occurrences beyond their previously known native ranges.

**Figure 7.**
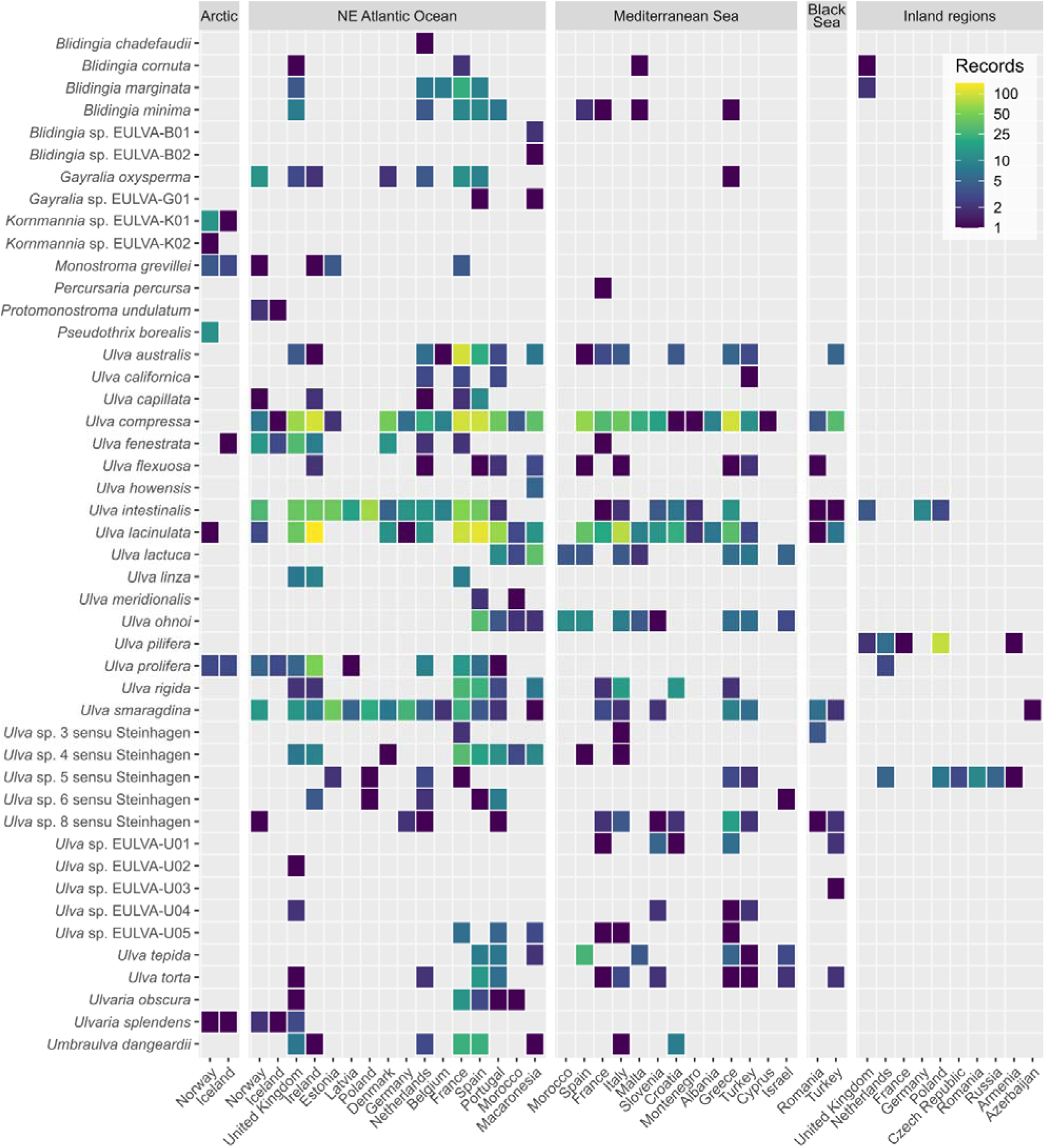
Heatmap showing the number of records for each of the 46 species across countries and regions. The boundary between the Arctic and NE Atlantic Ocean follows the marine ecoregions defined by Spalding et al. (2007). Inland regions comprise samples collected more than 20 km from the coast and include both freshwater habitats and landlocked saline lakes or brackish rivers. Colour intensity represents the number of records, with lighter colours indicating higher numbers of observations.

#### Ulvales – Ulvaceae

Ulva compressa was the most frequently collected species in the dataset (n = 856 records; Fig. 8A), followed by Ulva lacinulata (n = 739) and Ulva intestinalis (n = 427) (Supplementary Material I). Collectively, these three species were sampled across nearly the entire study area, ranging from northern Europe (Svalbard for U. lacinulata; Norway and Iceland for U. compressa and U. intestinalis) southwards to Morocco and the Canary Islands, and across the NE Atlantic, Mediterranean Sea, and Black Sea.

**Figure 8.**
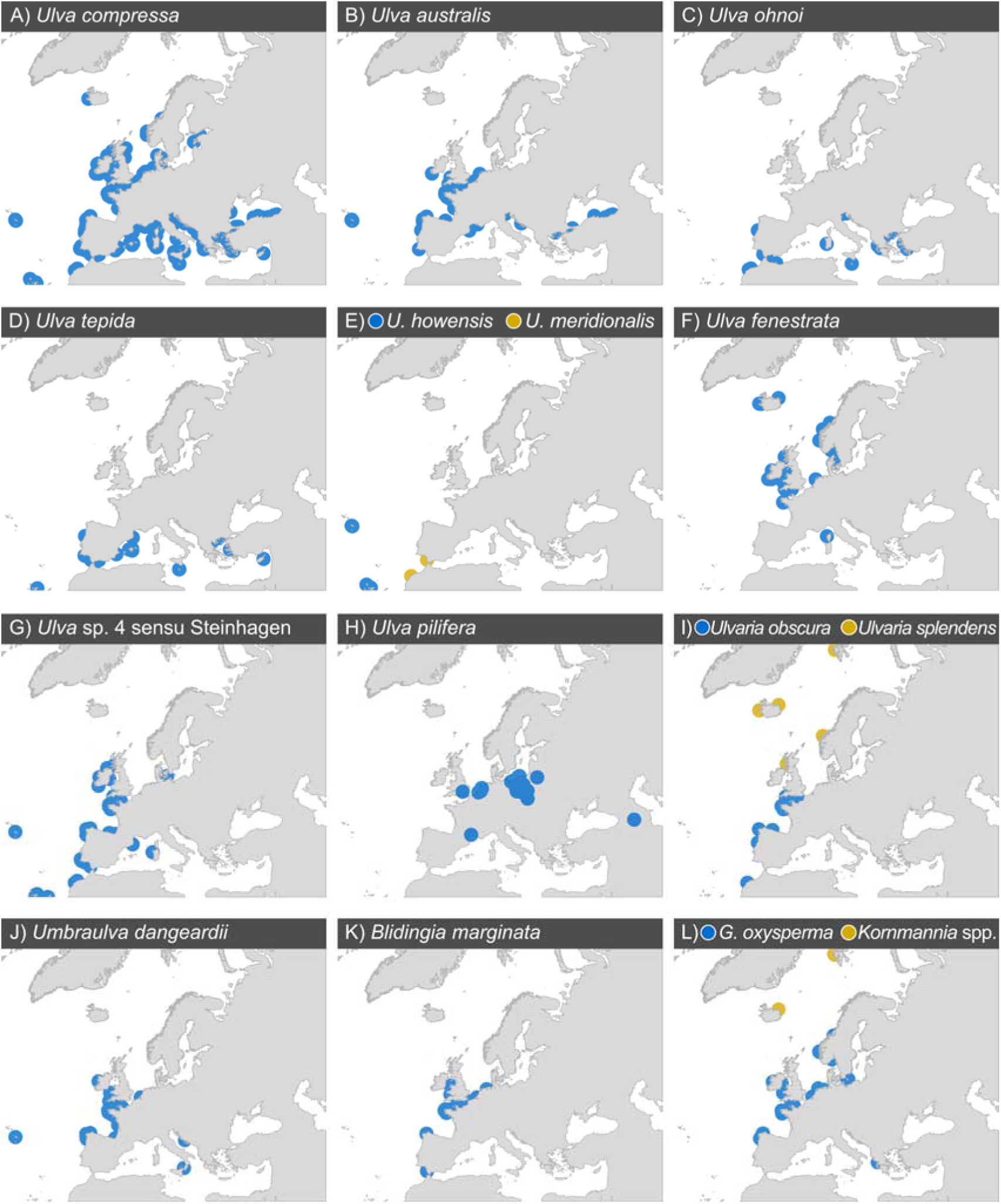
Distribution maps of 15 selected species of particular biogeographic interest recorded in the EULVA dataset. Occurrence records are displayed with 150 km buffers. (A) Ulva compressa was recorded throughout the study area, except in the Arctic. (B) The non-native Ulva australis has a broad distribution, including newly confirmed records for the UK, Croatia, and Macaronesia. (C) Ulva ohnoi exhibited a broader distribution than previously recorded. (D) Ulva tepida, previously known only from Israel, was detected at many additional locations. (E) Ulva howensis and Ulva meridionalis represent new records for the study area. (F) Ulva fenestrata and (G) Ulva sp. 4 sensu Steinhagen et al. (2023) were largely restricted to the NE Atlantic coast. (H) Ulva pilifera was associated with true freshwater habitats. (I) Ulvaria obscura was found in southern Europe, whereas Ulvaria splendens occurred from northern Europe to the Arctic. (J) Umbraulva dangeardii was recorded along the NE Atlantic coast and in the Mediterranean, often in subtidal habitats. (K) Blidingia marginata was exclusively found along the NE Atlantic coast. (L) Gayralia oxysperma occurred in temperate Europe, whereas Kornmannia species were restricted to northern Europe and the Arctic. See Supplementary Material I for distribution maps of all species.

Ulva australis, U. californica, U. lactuca, U. ohnoi, and U. tepida are considered non-indigenous species that were previously known to have been introduced to Europe and are native to the Pacific Ocean (Bartolo et al., 2022; van der Loos et al., 2024). Of these, U. australis was the most frequently collected species within this study (n = 169) and exhibited a broad distribution across the study area (Fig. 8B). It was collected from habitats varying from the rocky intertidal to pontoons, coastal lagoons, and maerl beds at depths of up to 22 m. Despite being recorded as early as 1984 in Europe (Gerbal et al., 1985; van der Loos et al., 2024), the present study represents the first record of U. australis from the United Kingdom, Croatia, and Macaronesia.

Both U. lactuca (n = 81) and U. ohnoi (n = 68) were predominantly found in the Mediterranean Sea and southern Europe, but neither were recorded on the Atlantic coast north of Spain. Ulva ohnoi was recorded for the first time in the Mediterranean (Israel) in 2002 and since then it has only been recorded in Italy, Portugal, Spain, and Tunisia (Coste et al., 2015; Krupnik et al., 2018; Miladi et al., 2018; Oca et al., 2019), and as fouling on hulls of ships (Flagella et al., 2010; Mineur et al., 2007). The EULVA dataset extends the known distribution of this species to include first records from Macaronesia, Malta, Slovenia, Greece, and Türkiye (Fig. 8C). The distribution of Ulva lactuca is historically more complex, as it was long considered native to European waters, but molecular analyses based on type material have shown it to be a tropical species. Specimens previously identified as Ulva lactuca in temperate and Arctic regions are now considered Ulva fenestrata (Hughey et al., 2019). This pattern is further supported by our genetic data, which indicate a characteristically northern distribution of Ulva fenestrata and a southern distribution of Ulva lactuca (Fig. 7).

Ulva tepida was previously only known from Israel, where it was first recorded in 2002 (Israel & Einav, 2017). Our dataset showed that this species is widespread (n = 56), with occurrences in the Mediterranean Sea (including Greece, Spain, and Türkiye) as well as the NE Atlantic Ocean (Spain, Portugal, and the Canary Islands) (Fig. 8D).

In contrast, U. californica was represented by only ten specimens but was nevertheless recorded from multiple locations across the NE Atlantic and Mediterranean Sea (Supplementary Material I). These are the first confirmed records of this species from France, Portugal, and Türkiye.

Ulva howensis and U. meridionalis were not previously known from Europe and may represent recent cryptic introductions. Ulva howensis was previously considered endemic to Australia (Kirkendale et al., 2013). Our four records from the Azores and Canary Islands therefore constitute the first records of this species outside Australia (Fig. 8E). Whether U. howensis is a previously overlooked native species or a recent introduction to the region remains unclear. Ulva meridionalis was originally described from Japan and has subsequently been confirmed from Southeast Asia (Monotilla & Tahil, 2023), Australia (Phillips et al., 2016), and the Western Atlantic (Melton & Lopez-Bautista, 2021). Our three records from Morocco and Spain represent the first records of this species within the study area (Fig. 8E).

Species predominantly recorded from the NE Atlantic included Ulva capillata, U. fenestrata, U. linza, and U. prolifera. Ulva capillata (n = 17), recently described from Sweden by Steinhagen et al. (2022), has previously been confirmed from Denmark, Norway, and the Netherlands. Our dataset indicates a dominant Atlantic distribution extending south to Spain and Portugal, where it was typically associated with sandy habitats and brackish riverbanks. Ulva fenestrata (n = 74) was largely restricted to northern Europe, with a single exception from Corsica in the Mediterranean Sea (Fig. 8F). Our sampling could not confirm the presence of Ulva linza and U. prolifera in the Mediterranean Sea (Supplementary Material I).

Similar to U. lactuca, comparisons between the current and historically reported distributions of Ulva rigida are complicated by its frequent misidentification as U. lacinulata. Molecularly confirmed records have so far been limited to Ireland, France, Spain, and Portugal (Hughey et al., 2022). Our dataset (n = 96) indicates that U. rigida is common along the Atlantic coast and throughout the Mediterranean, with records from the Azores, the United Kingdom, Italy, Croatia, and Greece (Supplementary Material I). The dataset also includes specimens collected from depths exceeding 20 m.

Three of the unnamed Ulva species recognised by Steinhagen et al. (2023) were relatively common: Ulva sp. 4 (n = 92), Ulva sp. 5 (n = 43), and Ulva sp. 8 (n = 35) (Supplementary Material I). Ulva sp. 4 was recorded predominantly along the NE Atlantic coast (Fig. 8G), while Ulva sp. 8 was reported mostly in the eastern Mediterranean and Black Sea, where it was frequently associated with saltwater creeks, coastal lagoons, saltmarshes, and brackish waters. Similarly, Ulva sp. 5 was primarily found in coastal lagoons and in brackish lakes and rivers, including landlocked saline lakes. In contrast, Ulva pilifera was recorded exclusively in freshwater habitats with no apparent saltwater influence (Fig. 8H).

The five additional unnamed Ulva species (EULVA-U01 to U05) were generally uncommon (Supplementary Material I). EULVA-U02 and EULVA-U03 were each represented by a single specimen, EULVA-U04 by seven specimens, EULVA-U01 by 32 specimens restricted to the Mediterranean and Black Sea, and EULVA-U05 by 17 specimens distributed across Macaronesia, the NE Atlantic, and the Mediterranean.

Finally, Ulvaria splendens (n = 8) displayed a distribution ranging from the Arctic to northern Europe, whereas U. obscura (n = 19) was restricted to temperate regions, extending from Brittany (France) to Morocco (Fig. 8I). Umbraulva dangeardii (n = 67) was recorded from both the NE Atlantic and Mediterranean and was often collected from subtidal habitats, down to 22 m depth (Fig. 8J).

#### Ulvales – Kornmanniaceae

Blidingia marginata (n = 52) and B. minima (n = 44) were the most common and widespread Blidingia species in our dataset. While B. marginata was only found on the NE Atlantic coast and adjacent seas (incl. the North Sea) (Fig. 8K), B. minima was also found in the Mediterranean Sea (Supplementary Material I). For Malta, this represents the first record of B. minima. Both species were found year-round and from mid-to high-intertidal areas.

Blidingia cornuta is a species recently described by Steinhagen et al. (2021) from Germany and was, until now, only known from locations in Germany along the Baltic Sea, Wadden Sea, and North Sea. Our sampling efforts resulted in five additional specimens from the United Kingdom, France, and Malta (Supplementary Material I). These included mostly locations in estuaries, with brackish waters and silt. This result shows that B. cornuta, although less common than B. marginata and B. minima, is more widespread than current records show.

The two unidentified Blidingia species (EULVA-B01 and B02) were recorded in Macaronesia (Canary Islands and Madeira) on rocky intertidal platforms (Supplementary Material I). Finally, a single specimen matching a reference sequence from the Canadian Atlantic coast identified as B. chadefaudii was found in the Netherlands (Supplementary Material I). None of the Blidingia species were recorded in the Arctic. On the contrary, the Kornmannia specimens (n = 15) were only found in Arctic regions (Fig. 8L).

#### Ulotrichales

The most frequently sampled member of the Ulotrichales was Gayralia oxysperma (n = 46), which showed a predominantly temperate distribution and was commonly found on mud-covered rocks in the high intertidal zone (Fig. 8L). Two specimens of an unidentified Gayralia species (EULVA-G01) were collected from the Azores and the Atlantic northern coast of Spain (Supplementary Material I).

Monostroma grevillei (n = 17) was collected from a range of habitats along the NE Atlantic coast and Arctic regions (but was not found south of Brittany, France), including harbours and as an epiphyte on other macroalgae species (Supplementary Material I). Pseudothrix borealis (n = 12) was found exclusively in Svalbard, consistent with its currently known Arctic distribution, which includes Russia and Alaska. However, these specimens represent the first record of the species from Svalbard. Finally, a single specimen of Percursaria percursa and three specimens of Protomonostroma undulatum were collected, the latter exclusively from northern Europe (Supplementary Material I).

## 4. Discussion

The EULVA dataset provides the most comprehensive, molecularly validated assessment of the diversity and geographical distribution of Ulvales and Ulotrichales across Europe and adjacent regions to date. By applying standardized DNA-based species identification to thousands of samples collected across the study area, we uncovered distribution patterns that were previously obscured by taxonomic uncertainty and morphological plasticity. Resolving these spatial patterns transforms how opportunistic macroalgae can be integrated into marine conservation and coastal management, for example in relation to green tides events that smother ecosystems (Ren et al., 2024; Wan et al., 2017) or the introduction of non-indigenous species. Historically, Ulva had the reputation of lacking biogeographic patterns, with species often reported to have broad distributions (Poole & Raven, 1997). This perception was particularly prevalent in the pre-DNA era, when European species names were frequently applied to specimens from other regions, leading to widespread reports of European species as globally distributed. Management of macroalgal blooms and coastal health currently still relies on these generic assumptions of species composition. By establishing a robust molecular baseline, our dataset enables the early detection of non-indigenous introductions, and provides a benchmark to track community shifts driven by eutrophication and climate change.

### 4.1 High-throughput Nanopore sequencing enables reliable species identification

Beyond resolving Ulva biogeography, the high sequencing success rate demonstrates that the workflow developed here is well-suited for large-scale, high-throughput taxonomic surveys. Nanopore-derived consensus sequences were virtually identical to Sanger sequences, confirming that the approach generates barcode-quality data for both ITS and tufA. We used a conservative approach throughout our study using consensus sequences based on at least 100 reads. However, our benchmarking analyses showed that consensus sequences generated from as few as 10-25 reads were identical to corresponding Sanger-derived sequences. This suggests that the throughput could potentially be increased further by lowering the required number of reads and multiplexing more samples per flow cell. Reducing the read threshold, however, also increases uncertainty in the resulting consensus sequences. The number of reads obtained per sample can indeed vary substantially, for example, due to differences in DNA quantity or quality, and is difficult to predict prior to sequencing. Moreover, very low read counts may result from contamination. Consensus sequences generated from low read numbers should therefore be interpreted with caution.

The ITS marker particularly benefited from the long-read sequencing strategy. Whereas mixed templates in Sanger sequencing often produce overlapping and noisy chromatograms that are difficult or impossible to interpret (Al-Shuhaib & Hashim, 2023; Shokralla et al., 2014), Nanopore sequencing resolves individual sequence variants into separate consensus sequences (Vierstraete & Braeckman, 2022). Consequently, samples containing multiple Ulva species or other eukaryotes can still yield accurate ITS barcodes without requiring additional cloning or repeated sequencing. Indeed, most samples contained multiple consensus sequences for both ITS and tufA amplicons, including the target Ulva sequence and sequences from associated organisms. These non-target sequences represent ‘bycatch’ from epiphytic, endophytic, or otherwise associated organisms, including the more commonly found fungi (ITS) and bacteria (tufA), but notably also small filamentous green algae such as Ulvella and Pseudendoclonium species (Nielsen et al., 2013). These understudied organisms are typically difficult to detect morphologically and even more challenging to sequence because of their intimate association with the host. As a result, these bycatch sequences may provide valuable information on the diverse communities inhabiting the thalli of foundational macroalgae, offering coastal managers and ecologists new opportunities to monitor and study cryptic host-associated biodiversity and the response of broader communities to environmental stress.

### 4.2 Taxonomic uncertainty

The identification of 46 species across more than 3,600 samples highlights both the extensive diversity of European Ulvales and Ulotrichales across Europe and adjacent regions and the continuing challenges associated with their taxonomy.

The persistence of taxonomic uncertainty is particularly evident within Ulva, where morphological similarity, phenotypic plasticity, and a history of taxonomic revisions have contributed to extensive nomenclatural confusion (Tran et al., 2022). Previous molecular taxonomic studies, particularly those incorporating sequences generated from type material, have already resolved the identity and nomenclature of several European species and substantially improved the taxonomic framework of the genus (Hughey et al., 2019, 2022; van der Loos et al., 2025). Nevertheless, several EULVA specimens could not be confidently assigned to described species because available reference sequences were either absent or associated with names that are no longer considered reliable. For example, Ulva sp. EULVA-U02 showed similarity to an ITS sequence labelled as “Enteromorpha intestinaloides”, but the former species name is now considered a synonym of U. compressa (van der Loos et al., 2025), whereas EULVA-U02 was clearly distinct from U. compressa type-derived sequences. Similarly, EULVA-U05 matched sequences labelled as Ulva clathrata, a name that is currently difficult to interpret because it has also been applied to specimens corresponding to Ulva sp. 4 sensu Steinhagen et al. (2023). However, all EULVA-U05 specimens originated from southern Europe, whereas Ulva sp. 4 has been reported across the NE Atlantic, which is consistent with the type locality of U. clathrata in Sweden (Hayden et al., 2003).

Additional cases further illustrate these taxonomic ambiguities. Ulva sp. 8 sensu Steinhagen et al. (2023) also matches sequences in GenBank identified as Ulva maeotica, a species originally described from the Black Sea region of Ukraine (Burova et al., 2011). This is consistent with our observations, as most specimens assigned to this entity originated from the eastern Mediterranean Sea and Black Sea, although the species appears to have a broader distribution. Likewise, Ulva sp. 5 sensu Steinhagen et al. (2023) may correspond to Ulva paradoxa based on GenBank ITS accessions (e.g., MT887231 and HM447561). Both examples demonstrate how potentially outdated or incorrect annotations in public sequence repositories can add to taxonomic uncertainty and complicate molecular species identification.

Even when type-derived sequences are available, taxonomic uncertainty may persist because of historical ambiguities surrounding the application of names. One example is a species represented by 100 specimens in our dataset, all collected from freshwater habitats. These specimens matched the holotype-derived tufA sequence of Ulva shanxiensis with 100% identity, but no ITS sequence is available from the type material of that species. The ITS sequences of the same specimens within our dataset matched GenBank accession HM447571, which was identified by Mareš et al. (2011) as Ulva flexuosa subsp. pilifera and cross-referenced with an rbcL sequence that the authors stated was derived from the type specimen. However, the specimen sequenced by Mareš et al. (2011) was in fact Kützing’s specimen of Enteromorpha corniculata Kützing 1849 (voucher L 0054611 in the Leiden herbarium), whereas the actual type specimen of Enteromorpha pilifera Kützing 1856 is represented by herbarium voucher L.4138900. Bliding (1963) transferred E. pilifera to Enteromorpha flexuosa subsp. pilifera (Kützing) Bliding and treated E. corniculata as a synonym. This entity was later transferred to Ulva pilifera (Kützing) Škaloud & Leliaert (2018), rendering Enteromorpha flexuosa subsp. pilifera a synonym of U. pilifera (Škaloud et al., 2018).

Ulva shanxiensis was described much later from a freshwater river in China (Chen et al., 2015) and therefore has no nomenclatural priority. However, because the true type specimen of U. pilifera has not yet been sequenced, it remains uncertain whether the freshwater lineage represented in our dataset should ultimately be referred to as Ulva pilifera or Ulva corniculata. Given that Bliding (1963) synonymised E. corniculata with E. flexuosa subsp. pilifera, now recognised as U. pilifera, we provisionally retain the name U. pilifera. This decision also promotes nomenclatural stability, as U. pilifera has long been used for freshwater Ulva and accurately reflects the ecological affinity of this lineage. This case illustrates the importance of integrating evidence from multiple genetic markers with careful nomenclatural evaluation.

### 4.3 The value of large-scale molecular biodiversity surveys

The extensive geographic coverage achieved through the EULVA network demonstrates the value of coordinated, continent-scale sampling initiatives for documenting marine biodiversity. The detection of numerous country-level first records and substantial range extensions highlights that the current distribution of many Ulva species remains incompletely documented, even in comparatively well-studied European coastal waters such as the United Kingdom and the Netherlands. These gaps are particularly evident for recently described, cryptic, and non-indigenous species, many of which have likely remained undetected due to a historical reliance on morphology-based identifications.

Distributional datasets of seaweeds are often compiled from independent local studies that differ in sampling effort, methodology, and taxonomic resolution, thereby limiting broad-scale comparisons. In contrast, the standardized workflow applied in this study enabled direct comparisons across habitats and regions, resulting in a dataset of unprecedented consistency and scale. Such collaborative efforts not only advance fundamental understanding of species distributions but also provide an essential baseline for detecting future range shifts associated with biological invasions and climate change. Repeated application of this framework on an even wider global scale will allow changes in the distribution of native and non-indigenous Ulva species to be monitored with substantially greater confidence than previously possible.

For opportunistic macroalgae like Ulva, high-resolution biogeographic mapping is directly relevant to coastal conservation, environmental policy, and the mitigation of massive algal blooms (“green tides”). Green tides inflict severe ecological and socio-economic damage by degrading benthic habitats, inducing hypoxia, and outcompeting native marine flora (Lyons et al., 2014; Smetacek & Zingone, 2013). Under environmental policy frameworks such as the European Union’s Water Framework Directive (WFD) and Marine Strategy Framework Directive (MSFD), macroalgal abundance and composition serve as key biological quality elements for assessing coastal ecological status. However, environmental monitoring routinely aggregates Ulva into a single generic indicator of eutrophication.

Our findings demonstrate that European Ulva assemblages are taxonomically diverse and composed of species with distinct geographic affinities, physiological traits, and invasion histories. Confounding native bloom-formers with invasive taxa or cryptic range-expanding species risks misinterpreting coastal ecosystem dynamics. Integrating scalable, molecularly validated baselines into routine environmental monitoring allows management authorities to accurately identify bloom-originating species, track the spread of non-indigenous macroalgae, and refine ecosystem-based conservation strategies across European seas.

## Acknowledgements

This work was initiated as a collaborative effort by members of the COST Action SeaWheat (CA20106). The authors thank the European Cooperation in Science and Technology (COST) for supporting this manuscript. In addition, we thank Marianne Ligthart, Bart van der Loos, and Mark Burton for collecting additional samples.

Olivier De Clerck was supported by the Research Foundation Flanders (Senior Research Project G015623N, ESFRI grants GOH3817N and I001621N), European Commission (EX-AQUA project, grant agreement No. 101159509). This study received Portuguese national funds from FCT - Foundation for Science and Technology through contracts UID/04326/2025, UID/PRR/04326/2025 and LA/P/0101/2020 (DOI:10.54499/LA/P/0101/2020). Céline Rebours’ contribution was financed by the Research Council of Norway (319577 SAFER-IMTA) and Møreforsking AS. Thomas Burel, Vincent Le Garrec, Anaëlle Bizien, and Mathieu Helias conducted part of the sampling for this study as part of the Rebent monitoring program (funded by the Brittany Regional Council and the DREAL) and the monitoring conducted through the European Water Framework Directive and the Marine Strategy Framework Directive. Work of the authors of the Università Politecnica delle Marche received financial support from the European Union - NextGenerationEU, under the National Recovery and Resilience Plan (NRRP) for Italy, Mission 4 Component 2 Investment 1.4 - Call for tender No. 3138 of 16 December 2021. The Slovenian component of the research was financially supported by the Slovenian Research and Innovation Agency (ARIS; grant no. P1-0237). Martina Orlando-Bonaca and Ana Lokovšek would like to thank Leon Lojze Zamuda and Domen Trkov for their valuable assistance during the fieldwork. Laurie C. Hofmann was supported by the European Union’s HORIZON EUROPE Research and Innovation Programme under grant agreement No 101084180, project NOVAFOODIES (Demonstration of innovative functional food production systems based on a sustainable value chain of marine and freshwater raw materials for conscientious European consumers). Funded by the European Union. The sampling of inland Ulva from Poland was supported by the National Science Centre, Poland, under grant OPUS 26 (2023/51/B/NZ8/01244). Hannah S. Earp, Charlotte Jennings, Pippa J. Moore and Heather Sugden were supported by an EU LIFE Programme grant (LIFE20/NAT/UK/00277).„Juliet Brodie and Rob Mrowicki were supported through the UK Darwin Tree of Life Project. Luisa Pia Düsedau and Inka Bartsch conducted the sampling on Svalbard as part of the project FACE-IT (The Future of Arctic Coastal Ecosystems – Identifying Transitions in Fjord Systems and Adjacent Coastal Areas). FACE-IT has received funding from the European Union’s Horizon 2020 research and innovation program under grant agreement No. 869154. The AWIPEV Arctic Research Base and the Kings Bay Marine Laboratory personnel are gratefully acknowledged for their support of the field campaign in Kongsfjorden. Sampling in Thuringia, Germany, was supported by the German Federal Ministry of Education and Research through the Thuringian Water Innovation Cluster (ThWIC; project VANAPLA, grant no. 03ZU1214BB). Sampling in Galicia, Spain, was supported by the Xunta de Galicia through NextGenerationEU and the European Maritime, Fisheries and Aquaculture Fund (EMFAF), within the Galicia Marine Science Programme under the Complementary Science Plans for Marine Science of the Ministerio de Ciencia e Innovación (Spain), included in the Recovery, Transformation and Resilience Plan (PRTR-C17.I1). Research conducted in Croatia was financially supported by the Croatian Science Foundation (HRZZ; grant No. IP-2019-04-6984). Ljiljana Iveša gratefully acknowledges Edi Gljušćić and Andrea Bilajac for their dedicated support during the fieldwork.

## Data Accessibility Statement

The ITS and tufA sequences, metadata, and photos of all 3,657 samples have been archived in the Barcode of Life Data Systems (BOLD; https://www.boldsystems.org) under project EULVA with accession numbers EULVA001-26 to EULVA3679-26. Physical voucher specimens prepared during this study were deposited in recognized herbaria within their respective countries of collection (Supplementary Table S2).

## Ethical statement

Our work follows all ethical and research integrity standards.

## Appendices

Supplementary Table S1. Published regional molecular studies of Ulva in Europe.

Supplementary Table S2. Metadata associated with the 3,657 collected samples.

Supplementary Table S3. Overview of the indexed primer pairs.

Supplementary Figure S1. Pairwise percentage identity matrix of Sanger-and Nanopore-derived tufA sequences.

Supplementary Figure S2. Phylogenetic tree containing all ITS sequences.

Supplementary Figure S3. Phylogenetic tree containing all tufA sequences.

Supplementary Materials I. Geographic distribution maps of all 46 species.

