## Supplementary Materials I for "Large-scale Nanopore sequencing resolves species diversity and distribution of European Ulva"

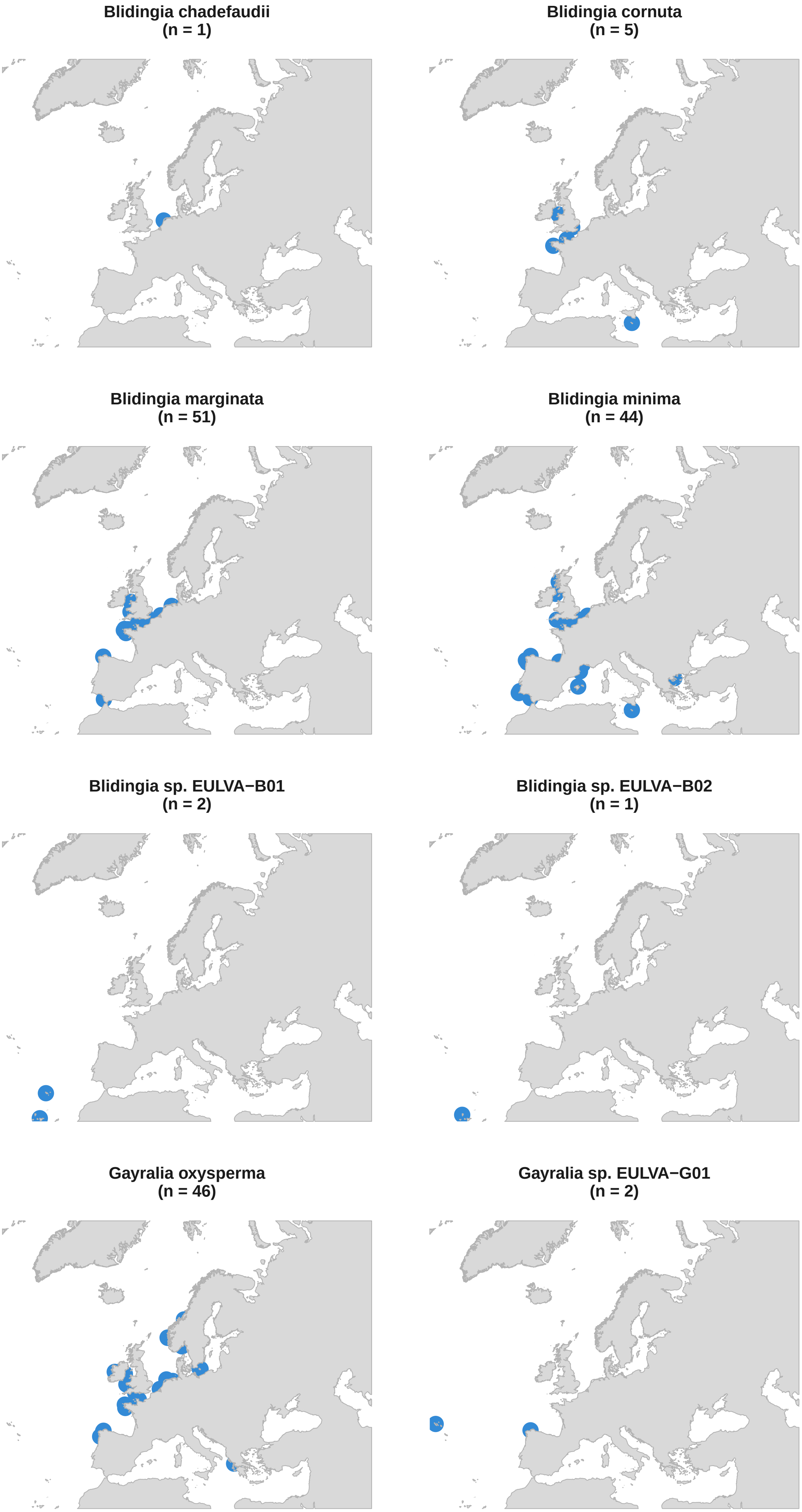
Blidingia chadefaudii
(n = 1)
Blidingia cornuta
(n = 5)
Blidingia marginata
(n = 51)
Blidingia minima
(n = 44)
Blidingia sp. EULVA−B01
(n = 2)
Blidingia sp. EULVA−B02
(n = 1)
Gayralia oxysperma
(n = 46)
Gayralia sp. EULVA−G01
(n = 2)

**Kornmannia sp. EULVA−K01**
**(n = 14)**

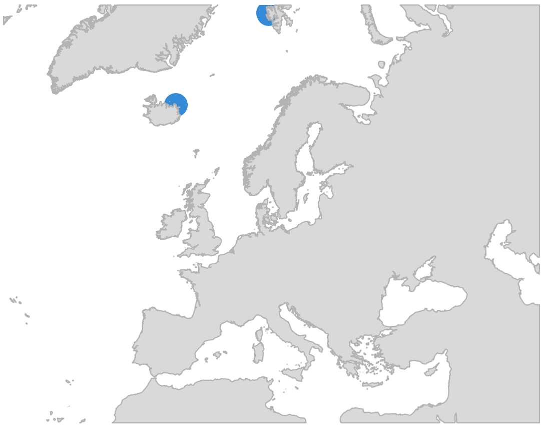

**Kornmannia sp. EULVA−K02**
**(n = 1)**

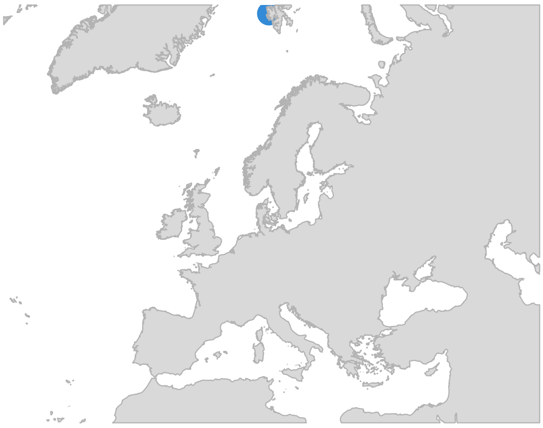

**Monostroma grevillei**
**(n = 18)**

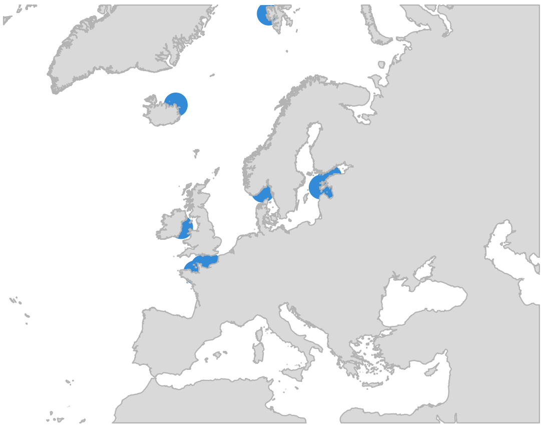

**Percursaria percursa**
**(n = 1)**

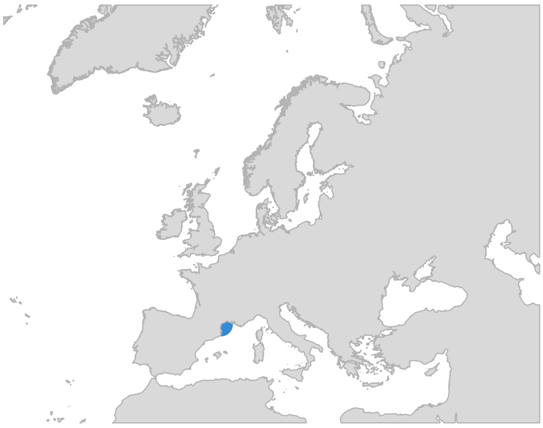

**Protomonostroma undulatum**
**(n = 3)**

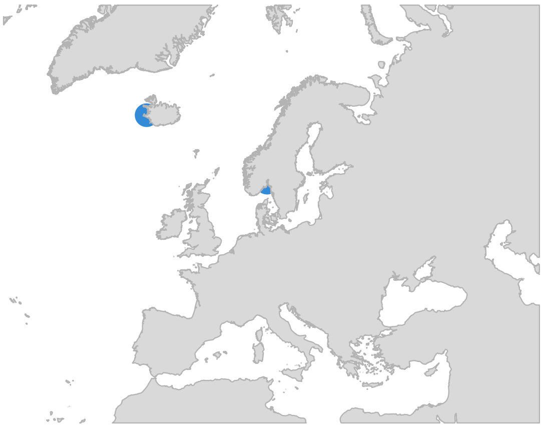

**Pseudothrix borealis**
**(n = 12)**

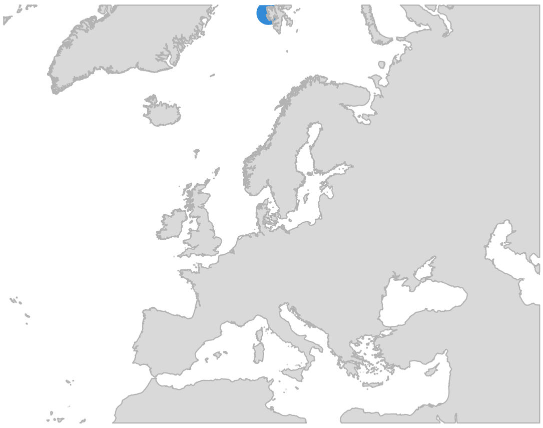

**Ulva australis**
**(n = 169)**

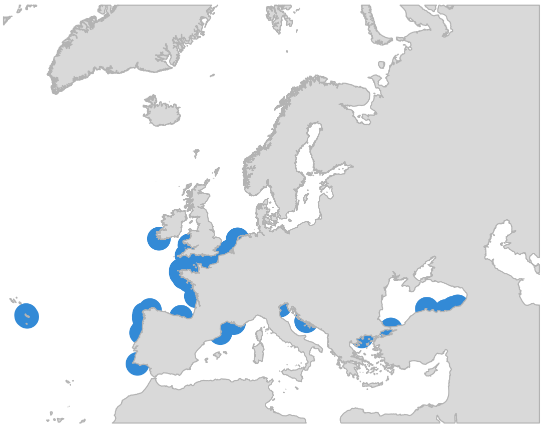

**Ulva californica**
**(n = 10)**

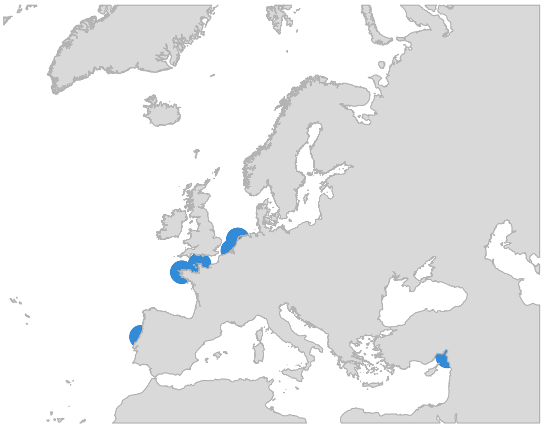

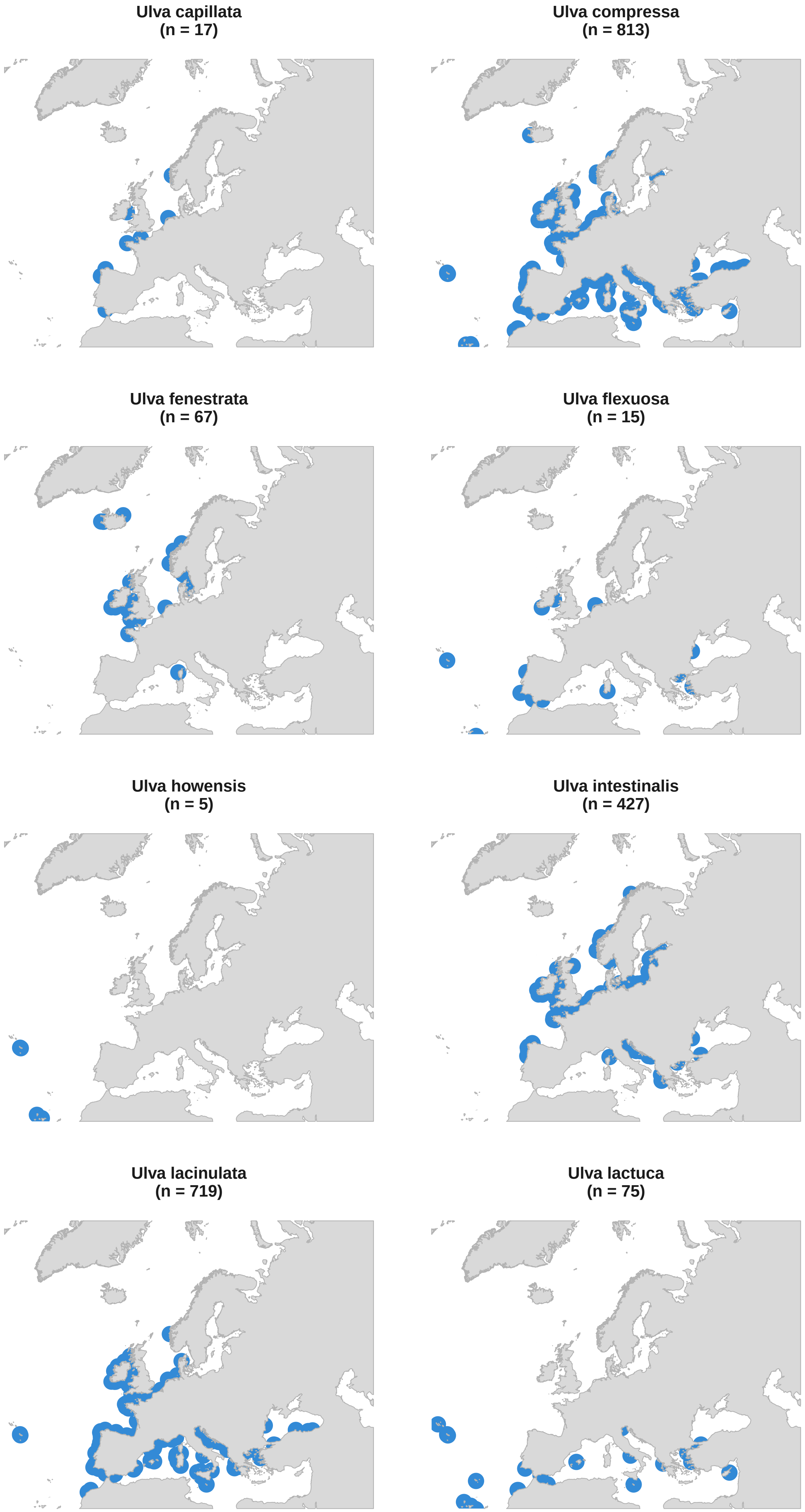
Ulva capillata
(n = 17)
Ulva compressa
(n = 813)
Ulva fenestrata
(n = 67)
Ulva flexuosa
(n = 15)
Ulva howensis
(n = 5)
Ulva intestinalis
(n = 427)
Ulva lacinulata
(n = 719)
Ulva lactuca
(n = 75)

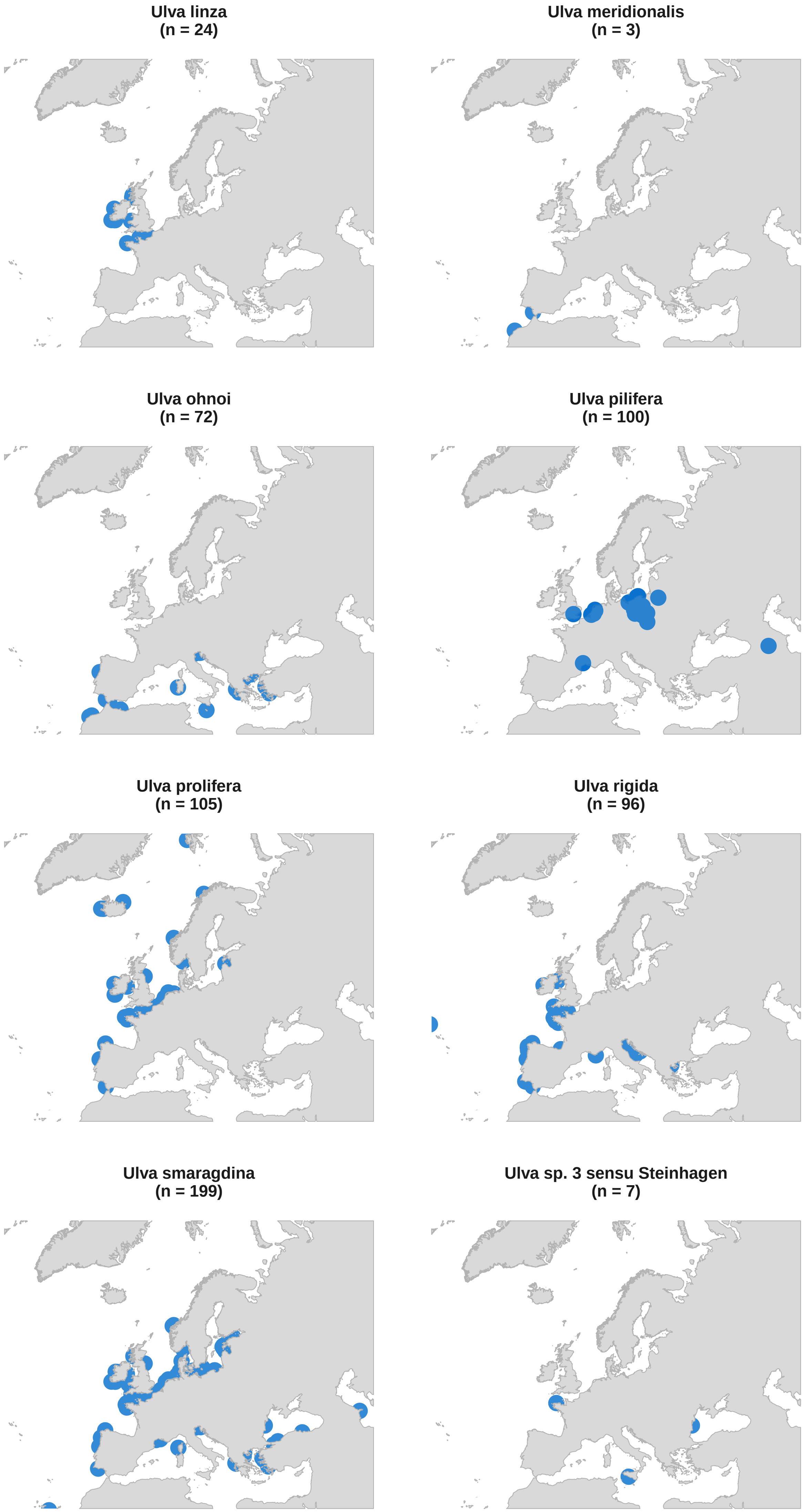
Ulva linza
(n = 24)
Ulva meridionalis
(n = 3)
Ulva ohnoi
(n = 72)
Ulva pilifera
(n = 100)
Ulva prolifera
(n = 105)
Ulva rigida
(n = 96)
Ulva smaragdina
(n = 199)
Ulva sp. 3 sensu Steinhagen
(n = 7)

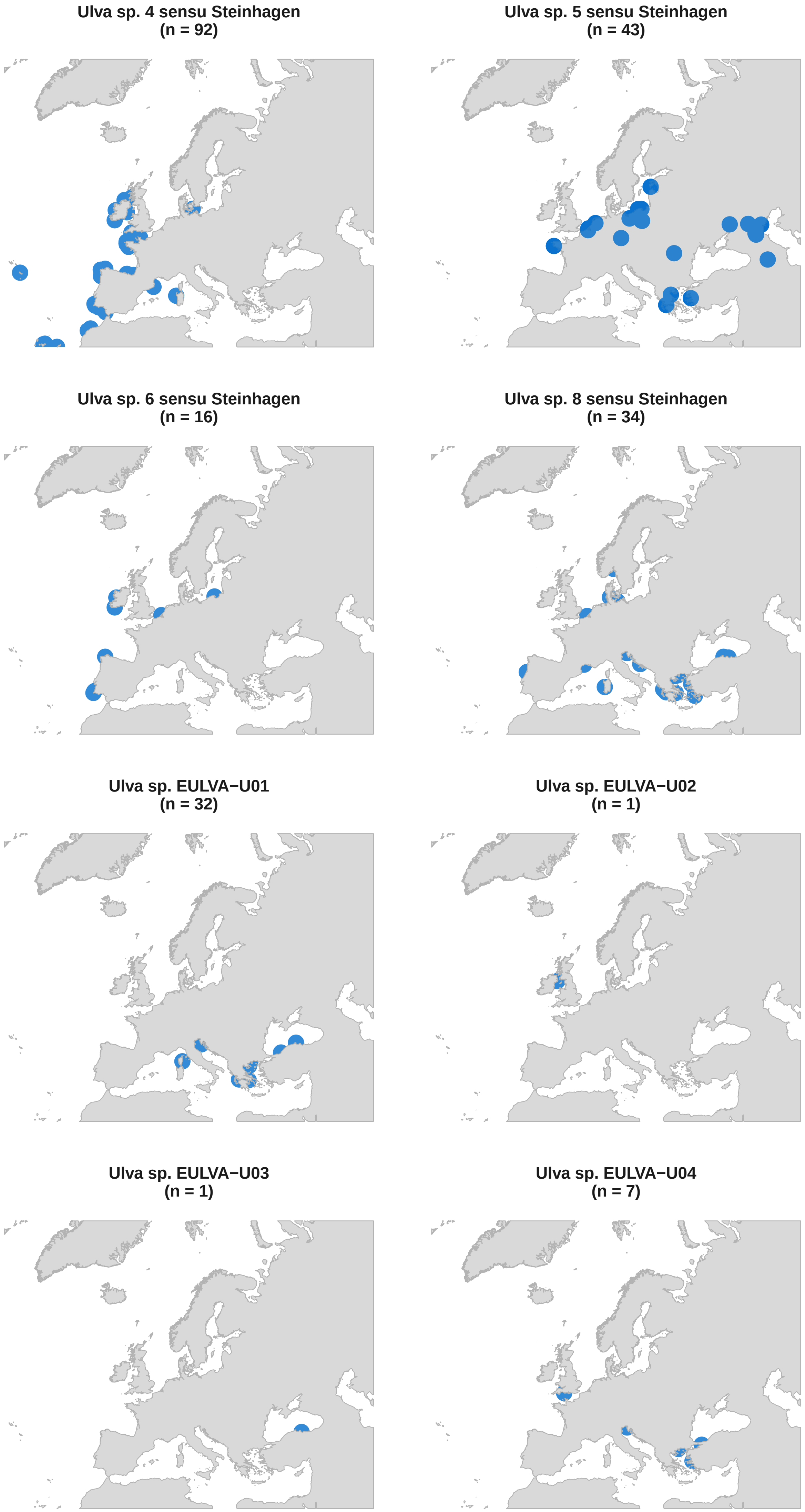
Ulva sp. 4 sensu Steinhagen
(n = 92)
Ulva sp. 5 sensu Steinhagen
(n = 43)
Ulva sp. 6 sensu Steinhagen
(n = 16)
Ulva sp. 8 sensu Steinhagen
(n = 34)
Ulva sp. EULVA−U01
(n = 32)
Ulva sp. EULVA−U02
(n = 1)
Ulva sp. EULVA−U03
(n = 1)
Ulva sp. EULVA−U04
(n = 7)

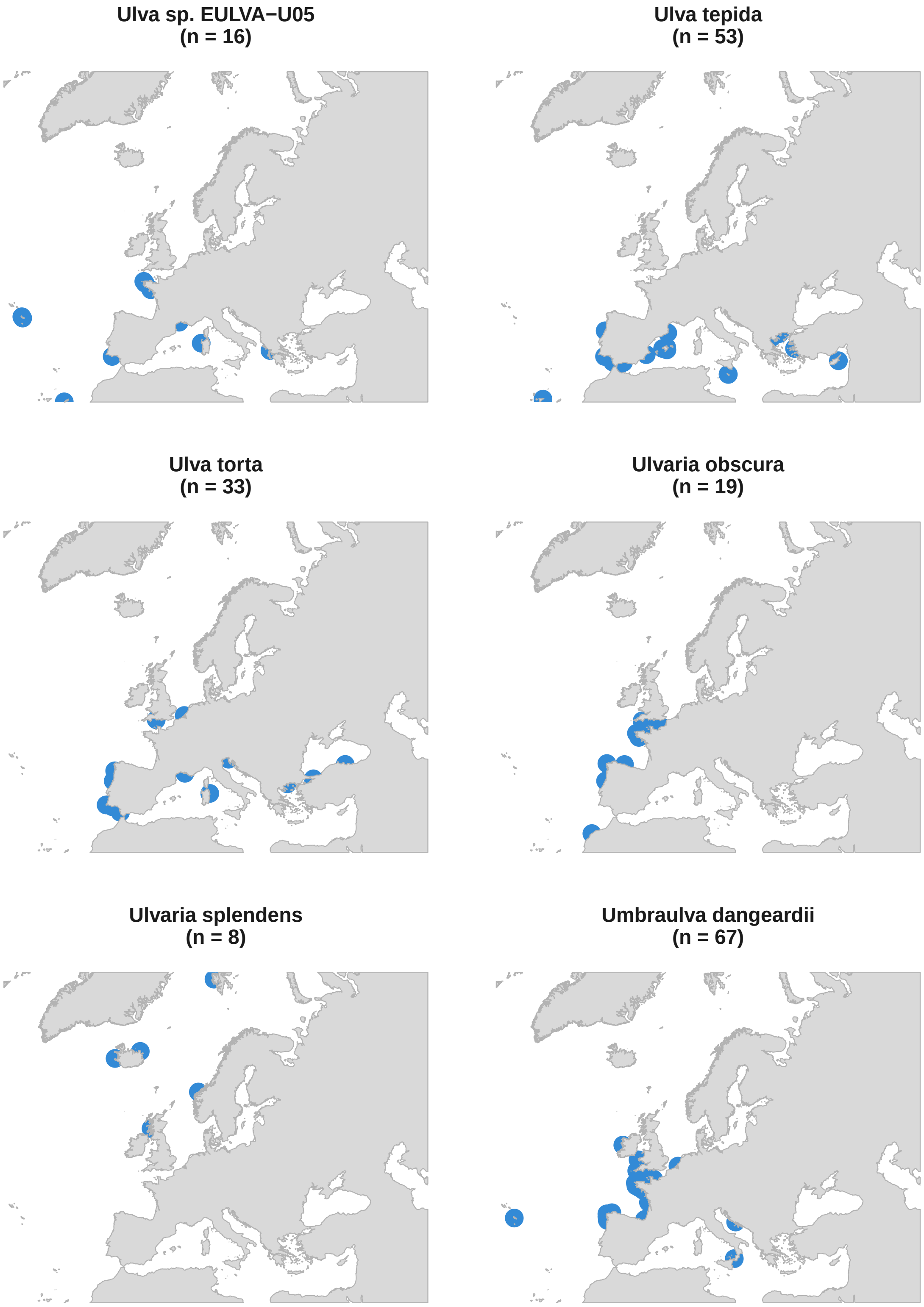
Ulva sp. EULVA−U05
(n = 16)
Ulva tepida
(n = 53)
Ulva torta
(n = 33)
Ulvaria obscura
(n = 19)
Ulvaria splendens
(n = 8)
Umbraulva dangeardii
(n = 67)
