## Supplementary material for "Large-scale Nanopore sequencing resolves species diversity and distribution of European Ulva": Table S1

**Supplementary Table S1.** Published regional molecular studies of *Ulva* in Europe.

| **Country or region** | **Habitat** | **DNA marker(s) used** | **Reference** | **Number of species recorded / main results** |
| --- | --- | --- | --- | --- |
| Italy: Lagoon of Venice and the Strait of Messina | Marine/transitional water | *tuf*A | (Armeli Minicante et al., 2024) | 10 species identified |
| Malta | Marine | ITS, *rbc*L, *tuf*A | (Bartolo et al., 2022) | 3 species identified |
| Norway: Bergen | Marine | *tuf*A | (Bringloe et al., 2019) | 2 species identified |
| Spain | Marine | ITS, *rbc*L | (Couceiro et al., 2011) | 1 species identified |
| France: Atlantic coast | Marine | *tuf*A | (Dartois et al., 2021) | 5 species identified |
| Portugal: Azores | Marine | *tuf*A | (Gabriel et al., 2024) | 4 species identified |
| Italy: Lagoon of Venice | Transitional water | *tuf*A | (Isocrono et al., 2026) | 1 species identified |
| France, UK | Marine | ITS, *rbc*L, *tuf*A | (Lamb et al., 2025) | 8 species identified |
| Denmark | Marine | *tuf*A | (Larsen-Ledet et al., 2025) | 5 species identified |
| Britain and Ireland | Marine | ITS, *rbc*L, *tuf*A | (Maggs et al., 2024) | Clarify nomenclature |
| Czech Republic, Finland, Germany, Hungary, Italy, Norway, Poland, Romania, Scotland and Sweden | Freshwater | ITS, *rbc*L | (Mareš et al., 2011) | 2 species identified |
| Italy | Marine | *tuf*A | (Miladi et al., 2018) | 10 species identified |
| Spain | Marine | *rbc*L | (Roth-Schulze et al., 2018) | 2 species identified |
| Poland | Freshwater | ITS, *rbc*L | (Rybak et al., 2014) | 2 species identified |
| Germany | Marine | *tuf*A | (Steinhagen et al., 2019) | 20 species identified |
| Denmark, Finland, Germany, Norway, Sweden | Marine | *tuf*A | (Steinhagen et al., 2023) | 20 species identified |
| Belgium, the Netherlands | Marine | *tuf*A | (van der Loos et al., 2025) | 22 species identified |
| Italy | Marine | *rbc*L, *tuf*A | (Wolf et al., 2012) | 6 species identified |
