## Supplementary material for "Large-scale Nanopore sequencing resolves species diversity and distribution of European Ulva": Figure S1

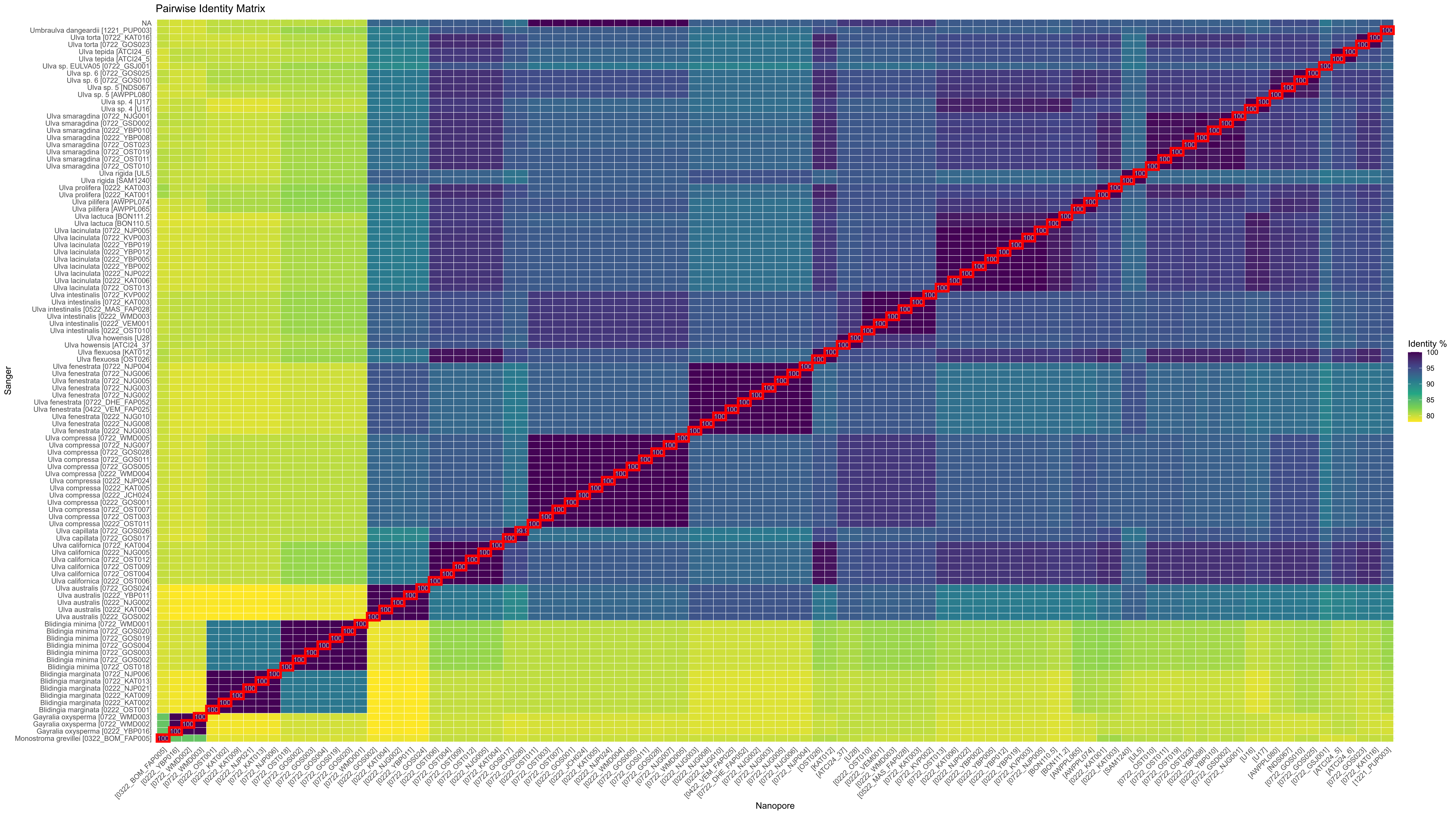

Pairwise Identity Matrix
NA
Umbraulva dangeardii [1221_PUP003]
Ulva torta [0722_KAT016]
Ulva torta [0722_GOS023]
Ulva tepida [ATCI24_6]
Ulva tepida [ATCI24_5]
Ulva sp. EULVA05 [0722_GSJ001]
Ulva sp. 6 [0722_GOS025]
Ulva sp. 6 [0722_GOS010]
Ulva sp. 5 [NDS067]
Ulva sp. 5 [AWPPL080]
Ulva sp. 4 [U17]
Ulva sp. 4 [U16]
Ulva smaragdina [0722_NJG001]
Ulva smaragdina [0722_GSD002]
Ulva smaragdina [0222_YBP010]
Ulva smaragdina [0222_YBP008]
Ulva smaragdina [0722_OST023]
Ulva smaragdina [0722_OST019]
Ulva smaragdina [0722_OST011]
Ulva smaragdina [0722_OST010]
Ulva rigida [UL5]
Ulva rigida [SAM1240]
Ulva prolifera [0222_KAT003]
Ulva prolifera [0222_KAT001]
Ulva pilifera [AWPPL074]
Ulva pilifera [AWPPL065]
Ulva lactuca [BON111.2]
Ulva lactuca [BON110.5]
Ulva lacinulata [0722_NJP005]
Ulva lacinulata [0722_KVP003]
Ulva lacinulata [0222_YBP019]
Ulva lacinulata [0222_YBP012]
Ulva lacinulata [0222_YBP005]
Ulva lacinulata [0222_YBP002]
Ulva lacinulata [0222_NJP022]
Ulva lacinulata [0222_KAT006]
Ulva lacinulata [0722_OST013]
Ulva intestinalis [0722_KVP002]
Ulva intestinalis [0722_KAT003]
Ulva intestinalis [0522_MAS_FAP028]
Ulva intestinalis [0222_WMD003]
Ulva intestinalis [0222_VEM001]
Ulva intestinalis [0222_OST010]
Ulva howensis [U28]
Ulva howensis [ATCI24_37]
Ulva flexuosa [KAT012]
Ulva flexuosa [OST026]
Ulva fenestrata [0722_NJP004]
Ulva fenestrata [0722_NJG006]
Ulva fenestrata [0722_NJG005]
Ulva fenestrata [0722_NJG003]
Ulva fenestrata [0722_NJG002]
Ulva fenestrata [0722_DHE_FAP052]
Ulva fenestrata [0422_VEM_FAP025]
Ulva fenestrata [0222_NJG010]
Ulva fenestrata [0222_NJG008]
Ulva fenestrata [0222_NJG003]
Ulva compressa [0722_WMD005]
Ulva compressa [0722_NJG007]
Ulva compressa [0722_GOS028]
Ulva compressa [0722_GOS011]
Ulva compressa [0722_GOS005]
Ulva compressa [0222_WMD004]
Ulva compressa [0222_NJP024]
Ulva compressa [0222_KAT005]
Ulva compressa [0222_JCH024]
Ulva compressa [0222_GOS001]
Ulva compressa [0722_OST007]
Ulva compressa [0722_OST003]
Ulva compressa [0222_OST011]
Ulva capillata [0722_GOS026]
Ulva capillata [0722_GOS017]
Ulva californica [0722_KAT004]
Ulva californica [0222_NJG005]
Ulva californica [0722_OST012]
Ulva californica [0722_OST009]
Ulva californica [0722_OST004]
Ulva californica [0222_OST006]
Ulva australis [0722_GOS024]
Ulva australis [0222_YBP011]
Ulva australis [0222_NJG002]
Ulva australis [0222_KAT004]
Ulva australis [0222_GOS002]
Blidingia minima [0722_WMD001]
Blidingia minima [0722_GOS020]
Blidingia minima [0722_GOS019]
Blidingia minima [0722_GOS004]
Blidingia minima [0722_GOS003]
Blidingia minima [0722_GOS002]
Blidingia minima [0722_OST018]
Blidingia marginata [0722_NJP006]
Blidingia marginata [0722_KAT013]
Blidingia marginata [0222_NJP021]
Blidingia marginata [0222_KAT009]
Blidingia marginata [0222_KAT002]
Blidingia marginata [0222_OST001]
Gayralia oxysperma [0722_WMD003]
Gayralia oxysperma [0722_WMD002]
Gayralia oxysperma [0222_YBP016]
Monostroma grevillei [0322_BOM_FAP005]
Sanger
Nanopore
[0322_BOM_FAP005]
[0222_YBP016]
[0722_WMD002]
[0722_WMD003]
[0222_OST001]
[0222_KAT002]
[0222_KAT009]
[0222_NJP021]
[0722_KAT013]
[0722_NJP006]
[0722_OST018]
[0722_GOS002]
[0722_GOS003]
[0722_GOS004]
[0722_GOS019]
[0722_GOS020]
[0722_WMD001]
[0222_GOS002]
[0222_KAT004]
[0222_NJG002]
[0222_YBP011]
[0722_GOS024]
[0222_OST006]
[0722_OST004]
[0722_OST009]
[0722_OST012]
[0222_NJG005]
[0722_KAT004]
[0722_GOS017]
[0722_GOS026]
[0222_OST011]
[0722_OST003]
[0722_OST007]
[0222_GOS001]
[0222_JCH024]
[0222_KAT005]
[0222_NJP024]
[0222_WMD004]
[0722_GOS005]
[0722_GOS011]
[0722_GOS028]
[0722_NJG007]
[0722_WMD005]
[0222_NJG003]
[0222_NJG008]
[0222_NJG010]
[0422_VEM_FAP025]
[0722_DHE_FAP052]
[0722_NJG002]
[0722_NJG003]
[0722_NJG005]
[0722_NJG006]
[0722_NJP004]
[OST026]
[KAT012]
[ATCI24_37]
[U28]
[0222_OST010]
[0222_VEM001]
[0222_WMD003]
[0522_MAS_FAP028]
[0722_KAT003]
[0722_KVP002]
[0722_OST013]
[0222_KAT006]
[0222_NJP022]
[0222_YBP002]
[0222_YBP005]
[0222_YBP012]
[0222_YBP019]
[0722_KVP003]
[0722_NJP005]
[BON110.5]
[BON111.2]
[AWPPL065]
[AWPPL074]
[0222_KAT001]
[0222_KAT003]
[SAM1240]
[UL5]
[0722_OST010]
[0722_OST011]
[0722_OST019]
[0722_OST023]
[0222_YBP008]
[0222_YBP010]
[0722_GSD002]
[0722_NJG001]
[U16]
[U17]
[AWPPL080]
[NDS067]
[0722_GOS010]
[0722_GOS025]
[0722_GSJ001]
[ATCI24_5]
[ATCI24_6]
[0722_GOS023]
[0722_KAT016]
[1221_PUP003]
Identity %
100
95
90
85
80
