## Supplementary figures and images for "Large-scale Nanopore sequencing resolves species diversity and distribution of European Ulva"

### Figure S2

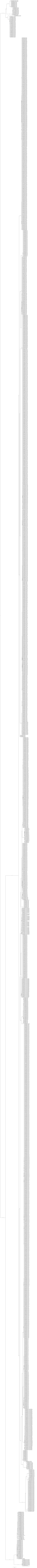

### Figure S3

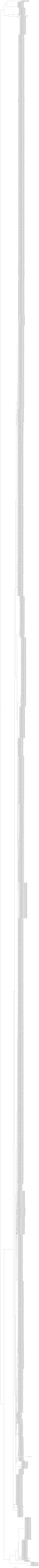
